# Enterotoxigenic *Escherichia coli* heat-labile toxin drives enteropathic changes in small intestinal epithelia

**DOI:** 10.1101/2022.08.24.504189

**Authors:** Alaullah Sheikh, Brunda Tumala, Tim J. Vickers, John C. Martin, Bruce A. Rosa, Subrata Sabui, Supratim Basu, Rita D. Simoes, Makedonka Mitreva, Chad Storer, Erik Tyksen, Richard D. Head, Wandy Beatty, Hamid M. Said, James M. Fleckenstein

## Abstract

Enterotoxigenic *E. coli* (ETEC), produce heat-labile (LT) and/or heat-stable (ST) enterotoxins, and are a common cause of diarrhea in children of resource-poor regions. ETEC have also been linked repeatedly to poorly understood sequelae including enteropathy, malnutrition, and growth impairment. While the cellular actions of ETEC enterotoxins leading to diarrhea are well-established, their potential contribution to subsequent pathology is unclear. LT stimulates cellular cAMP production to activate protein kinase A (PKA) which phosphorylates cellular ion channels that drive export of salt and water into the intestinal lumen resulting in diarrhea. However, as PKA exhibits broad kinase activity and its activated catalytic subunits modulate transcription of many genes, we interrogated the transcriptional profiles of LT-treated small intestinal epithelia. These studies demonstrated toxin-induced changes in hundreds of genes including those required for biogenesis and function of the brush border, the major site absorption of nutrients, and suppression of a key transcription factors, HNF4 and SMAD4, critical to differentiation of intestinal epithelia. Accordingly, LT treatment of intestinal epithelial cells significantly disrupted the absorptive microvillus architecture and altered transport of essential nutrients. In addition, challenge of neonatal mice with LT-producing ETEC recapitulated the architectural derangement of the brush border while maternal vaccination with LT prevented brush border disruption in ETEC-challenged neonatal mice. Finally, mice repeatedly challenged with toxigenic ETEC exhibited impaired growth recapitulating the multiplicative impact of recurring ETEC infections in children. These findings highlight impacts of ETEC enterotoxins beyond acute diarrheal illness and may inform approaches to mitigate and prevent major sequelae including malnutrition that impact millions of young children.

## Introduction

Infectious diarrhea remains a leading cause of death and morbidity among young children in low-middle income countries where access to clean water and sanitation remain in short supply^1^. Enterotoxigenic *E. coli* (ETEC), initially discovered as a cause of severe, cholera-like illness^2^, are one of the most common pathogens associated with moderate-severe diarrhea among children under the age of five years^3,4^, and are perennially the most common cause of diarrhea in travelers^5^ to endemic regions where these organisms are thought to account for hundreds of millions of cases of diarrheal illness each year^6^.

Importantly, ETEC infections have been linked to non-diarrheal sequelae including “environmental enteric dysfunction (EED),” a condition characterized by impaired nutrient absorption, impaired growth^7,8^ and malnutrition,^9,10^ adding significantly to the morbidity as well as deaths from diarrhea and other infections^11^. The risk of stunting multiplies with each episode of diarrheal illness in children under the age of two years^12^, a period during which children residing in impoverished areas commonly sustain multiple ETEC infections^8^. However, the molecular pathogenesis underlying the intestinal changes associated with EED, and the contribution of individual pathogens including ETEC remain poorly understood.

Similarly, toxin-producing *E. coli* have also been repeatedly identified in patients with tropical sprue^13–15^, a condition classically described in adults residing for extended periods of time in areas where ETEC diarrheal disease is common. Like EED, tropical sprue is associated with changes to the small intestinal villous architecture, including ultrastructural alteration of the epithelial brush border formed by the microvilli^16,17^, nutrient malabsorption, and wasting.

The basic molecular mechanisms underpinning acute watery diarrhea caused by ETEC are well-established^18^. ETEC produce heat-labile (LT) and/or heat-stable (ST) enterotoxins that activate production of cAMP and cGMP second messengers respectively, leading to activation of cellular kinases that in turn modulate the activity of sodium and chloride channels in apical membrane of intestinal epithelial cells to promote net efflux of salt and water into the intestinal lumen resulting in watery diarrhea.

LT and cholera toxin (CT) share ~85 % amino acid identity, and both toxins exert their major effects on the cell through the ADP-ribosylation of the alpha subunit of Gs (Gsα), a stimulatory intracellular guanine nucleotide-binding protein. Inhibition of Gsα GTPase activity leads to constitutive activation of adenylate cyclase and increased production of intracellular cAMP^19^.

In its central role as a second messenger, cAMP governs a diverse array of cellular processes^20^ and modulates transcription of multiple genes through a number of cAMP-responsive transcriptional activators and repressors^21^. cAMP activates protein kinase A (PKA), a heterotetramer, by liberating its two regulatory subunits from the catalytic subunits which are then free to phosphorylate a wide variety of cytoplasmic and nuclear protein substrates^22^. PKA largely regulates transcription by phosphorylation of transcription factors including the cyclic AMP response element binding protein (CREB) and the cAMP-response element modulator (CREM) which bind cAMP response elements (CRE) in the promoter regions of target genes^21–23^.

Notably, cholera toxin (CT), LT, and dibutyryl-cyclic AMP all induce hypersecretion and impact the architecture of gastrointestinal epithelia in rodent small intestine ^24^, while small intestinal biopsies of patients with acute cholera exhibit marked changes in the ultrastructure of the intestinal brush border, the major absorptive surface in the small intestine^25,26^, including shortening and disruption of the microvilli. Consistent with these observations, studies of young children less than two years of age in Bangladesh have specifically associated LT-producing ETEC with undernutrition^27^, suggesting that heat-labile toxin may exert effects on intestinal mucosa that extend beyond acute diarrheal illness.

Here we demonstrate that in addition to the canonical effects of LT on cellular export of salt and water into the intestinal lumen, this toxin impacts multiple genes involved in formation of microvilli, resulting in marked alteration of the architecture of the intestinal brush border, the major site of nutrient absorption in the small intestine. These effects are compounded by alteration of solute transporters within the brush border epithelia, potentially disrupting the absorption of multiple essential nutrients.

## Results

### Heat-labile toxin markedly alters the transcriptomes of intestinal epithelial cells

Commensurate with the importance of cAMP as a second messenger, we found that compared to either untreated controls or cells treated with a catalytically inactive (E211K) mutant of LT, wild type LT holotoxin substantially modulated transcription of many genes in intestinal epithelial cells. In RNAseq studies of polarized Caco-2 intestinal epithelial cells we found that 3,832 genes were significantly (p≤10^−5^) upregulated and 3687 downregulated in response to LT, while the inactive toxin failed to induce significant changes in the transcriptome (figure 1a). However, Caco-2 cells are derived from distant metastases of a colon cancer tumor in which transcriptomes would likely be altered relative to untransformed intestinal epithelia^28,29^, and cAMP signaling is known to be aberrant in some transformed cells^23^. Therefore, to examine a more physiologically relevant target, we next examined the impact of LT on differentiated small intestinal enteroids. Here, we found that far fewer genes were differentially expressed (≤10^−5^) with 746 significantly upregulated, and 561 downregulated in response to intoxication with LT (figure 1b). Notably, however we found substantial statistically significant overlap in genes significantly modulated in Caco-2 cells and enteroids (supplemental table 3, <u>supplemental dataset 1</u>) with the transcription of hundreds of genes significantly up- or down-regulated in both groups. Gene ontology enrichment analysis as well as ontology-independent investigation of genes (CompBio, supplemental figure 1) modulated by the toxin highlighted multiple cellular components associated with both the development and function of the absorptive surface of the small intestine (<u>supplemental dataset 2</u>).

**Figure 1.**
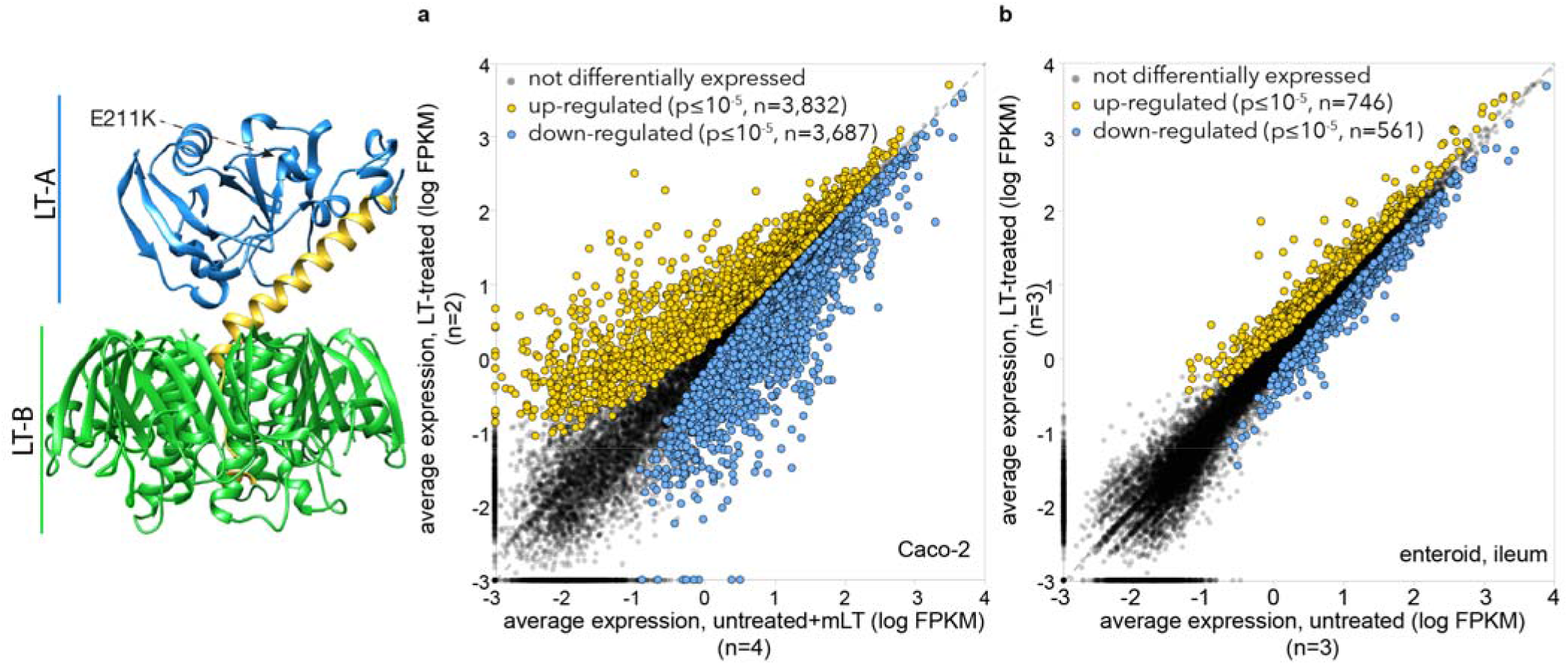
Heat-labile toxin modulates expression of multiple genes in intestinal epithelia. Model at left depicts the *E. coli* heat-labile toxin^92^ based on PDB structure <u>1LTS</u> with the A1 subunit in blue the A2 region of in yellow and pentameric B subunit in green. The E211K mutation of mLT is in the active site of the A1 subunit. **A.** Scatterplot of RNAseq data right depicts differential expression profiles of Caco-2 cells following exposure to heat-labile toxin (n=2) relative to untreated cells (n=2) and cells treated with the biologically inactive mLT (n=2). (Because expression profiles of untreated and mLT-treated cells were virtually identical, their combined expression profiles totaling n=4 replicates are compared here to LT-treated cells). **B.** RNAseq data from polarized small intestinal ileal enteroids treated with LT (n=3) compared to control untreated (n=3) cells.

### Heat-labile toxin impairs development of small intestinal microvilli

Small intestinal enterocytes are each covered with hundreds of microvilli, complex structures comprised of a central core of actin filaments within protrusions of the plasma membrane. Collectively, the luminal surface of the intestine formed by these microvilli, known as the brush border, represents the major absorptive surface of the gastrointestinal tract. Three major classes of proteins are required for the biogenesis of microvilli^30^ (figure 2a). These include (1) proteins such as villin, epsin, plastin, and EPS8 that bundle parallel clusters of actin filaments; BAIP2L1 (IRTKS) responsible for recruiting the EPS bundling protein to the tips of microvilli^31^; (2) ezrin, myo1a^32^, myo6 that link the actin cytoskeleton with the plasma membrane; and (3) protocadherin molecules CDHR2 and CDHR5 engaged in extracellular heterotypic complexes between the tips of the microvilli^30^ that are stabilized by a tripartite complex of MYO7B, ANKS4B^33^, and USH1C^33^. Interrogation of transcriptional profiles indicated that the transcription of each of these classes of genes was significantly altered following exposure to wild type heat-labile toxin (figure 2b). Similarly, RT-PCR confirmed decreased expression of multiple genes involved in microvilli biogenesis, including VIL1 encoding villin (figure 3a), and we were able to demonstrate that production of villin was depressed in polarized small intestinal enteroids (figure 3b supplemental figure 2). In addition, TEM images of polarized small intestinal enteroids exposed to heat-labile toxin demonstrated significantly shortened and disorganized microvillus structures on the apical surface of enterocytes (figure 3c).

**Figure 2.**
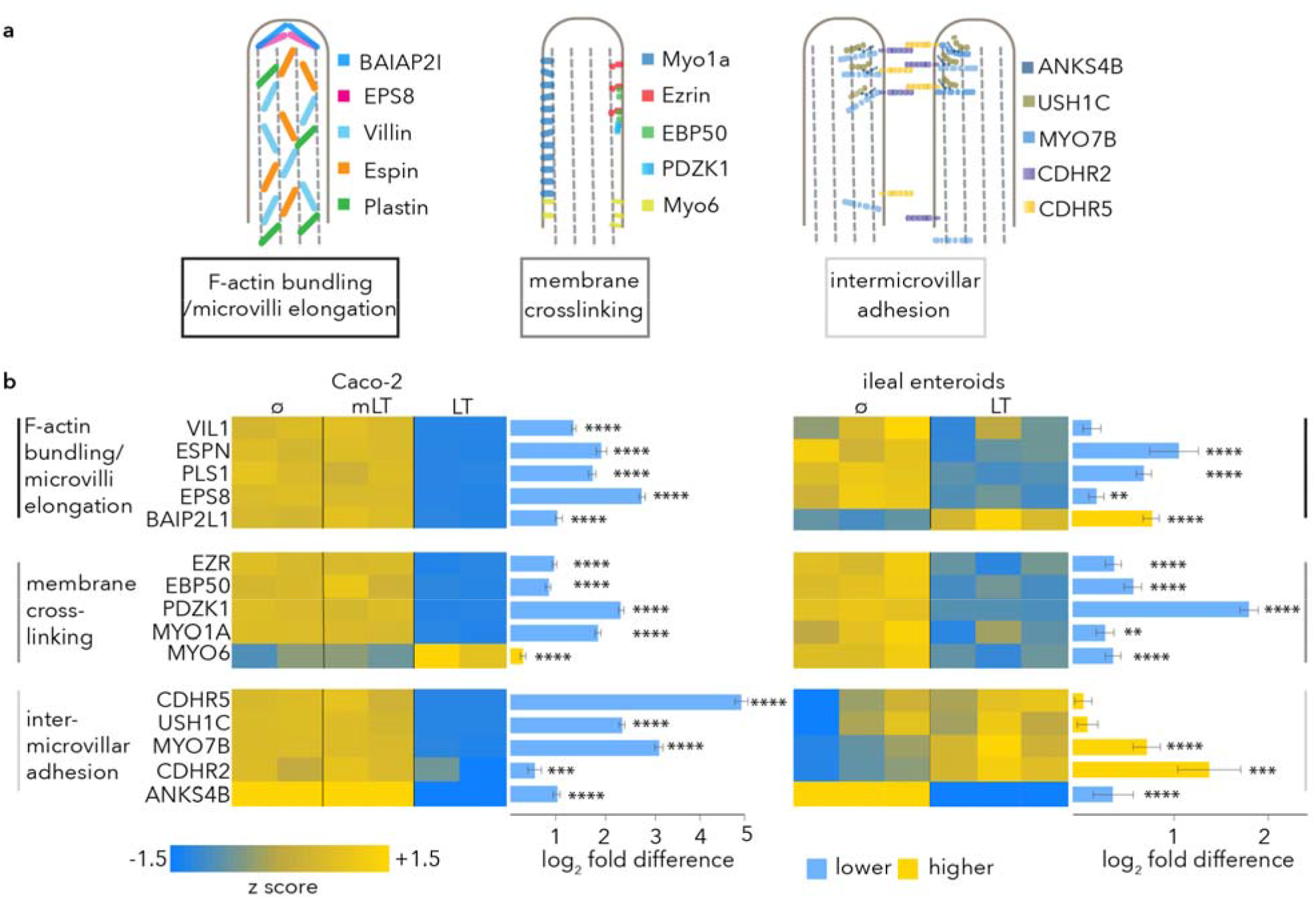
Heat labile toxin modulates multiple genes involved in microvillus assembly. **a.** diagram at top (adapted from reference^30^) depicts molecules involved in key elements of microvillus development. **b.** heat maps of RNAseq data obtained following treatment of Caco-2 intestinal cells (left) with mLT (n=2), or LT (n=2) relative to untreated cells (n=2); and ileal enteroids (right) treated with LT (n=3) relative to control untreated cells (n=3). RNAseq fold change data comparisons were made with DESeq2^93^. Bars indicate absolute fold change values + SE. *p≤0.05, ** p≤0.01, *** p≤0.001, **** p≤10^−4^, *****p≤10^−5^.

**Figure 3.**
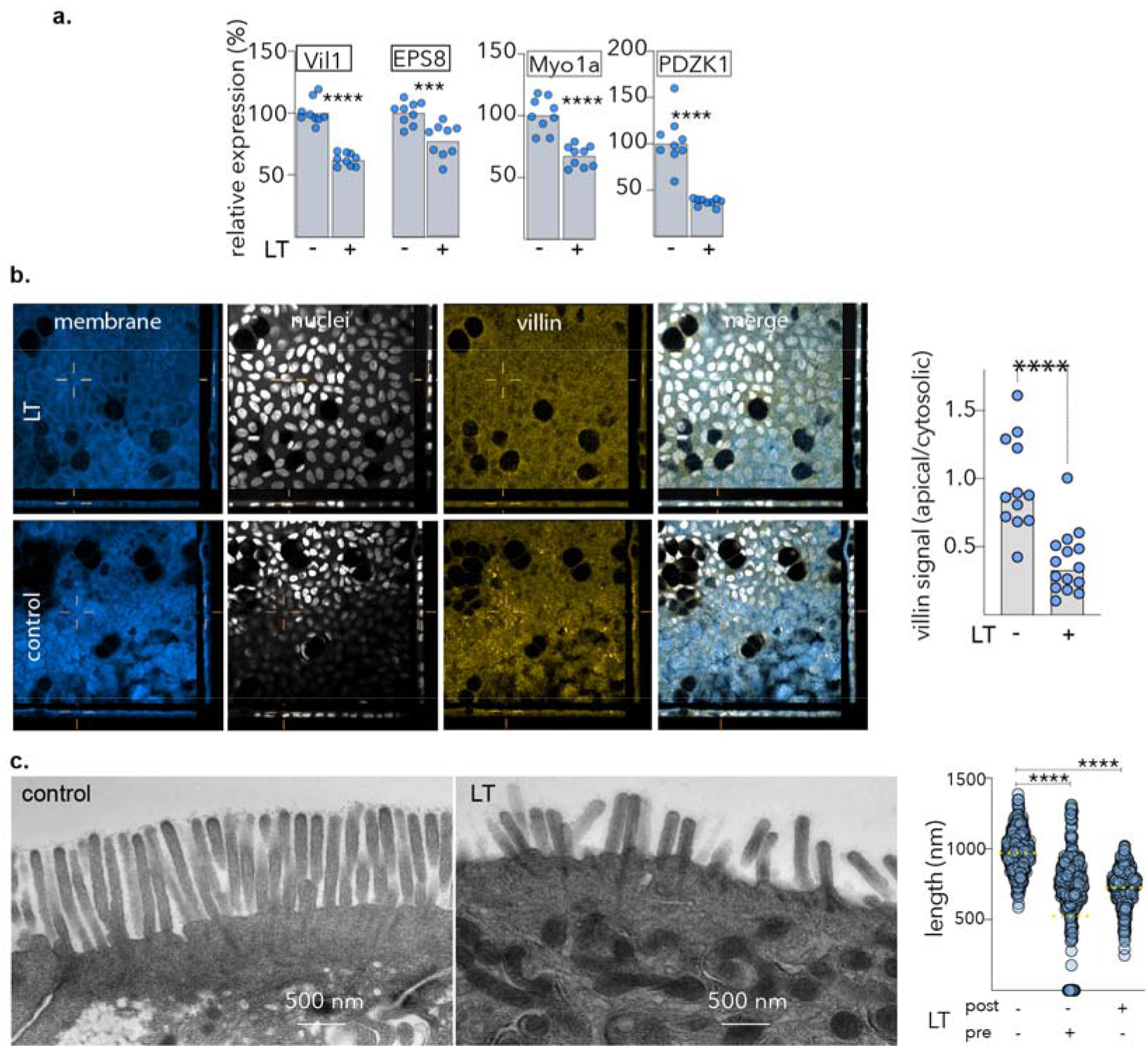
Heat labile toxin impairs effective formation of small intestinal microvilli. **a.** quantitative RT-PCR data for selected microvillus genes comparing untreated (-) and LT-treated (+) ileal epithelial cells (enteroid line 235D). **b.** Villin production is suppressed in small intestinal enteroids following treatment (t+18h) with LT (100 μg/ml). Shown are representative confocal images obtained showing membrane (CellMask, blue), nuclei (white), villin (gold), and merged image. Graph at right shows apical villin geometric mean fluorescence intensity data relative to the corresponding cytoplasmic signal. Each symbol (n=12)represents a unique region of interest. ****p<0.0001, Mann-Whitney 2 tailed testing. **c.** TEM images of small intestinal microvilli following treatment LT (right) compared to control untreated cells (left). Graph at right shows length of microvilli when enteroids are treated before (pre) and after (post) differentiation on polarized ileal cells ****<0.0001 by ANOVA (Kruskal-Wallis, nonparametric testing).

### Heat-labile toxin modulates the transcription of multiple brush border nutrient transport genes

The human solute carrier (SLC) gene superfamily is comprised of more than 50 gene families thought to encode more than 300 functional transporters^34^. Many of the SLC proteins are enriched in the small intestinal brush border where they transport critical nutrients including amino acids, oligopeptides, sugars, and vitamins. We found that transcription of many SLC genes was altered in Caco-2 cells as well as small intestinal enteroids (figure 4a). These included transporters for Zinc, known both to be deficient in children with enteropathy^35^, and a micronutrient critical for intestinal homeostasis. Likewise transcription of SLC19A3 encoding the principal SLC responsible for uptake of the water-soluble B vitamin thiamine (vitamin B1)^36^ by differentiated intestinal epithelial cells lining the surface of intestinal villi of the proximal small intestine^37,38^(figure 4b) was repressed as was production of the corresponding protein (supplemental figure 3a). Moreover, we found that LT treatment of human small intestinal organoids also interfered with transcription of the cis regulatory element specificity protein 1 (SP-1) previously shown to govern the transcription of SLC19A3^39–41^ (figure 4c, supplemental figure 3b). Finally, we found that thiamine transport was significantly depressed following exposure to LT (figure 4d) providing additional evidence that ETEC can impair transport of critical nutrients.

**Figure 4.**
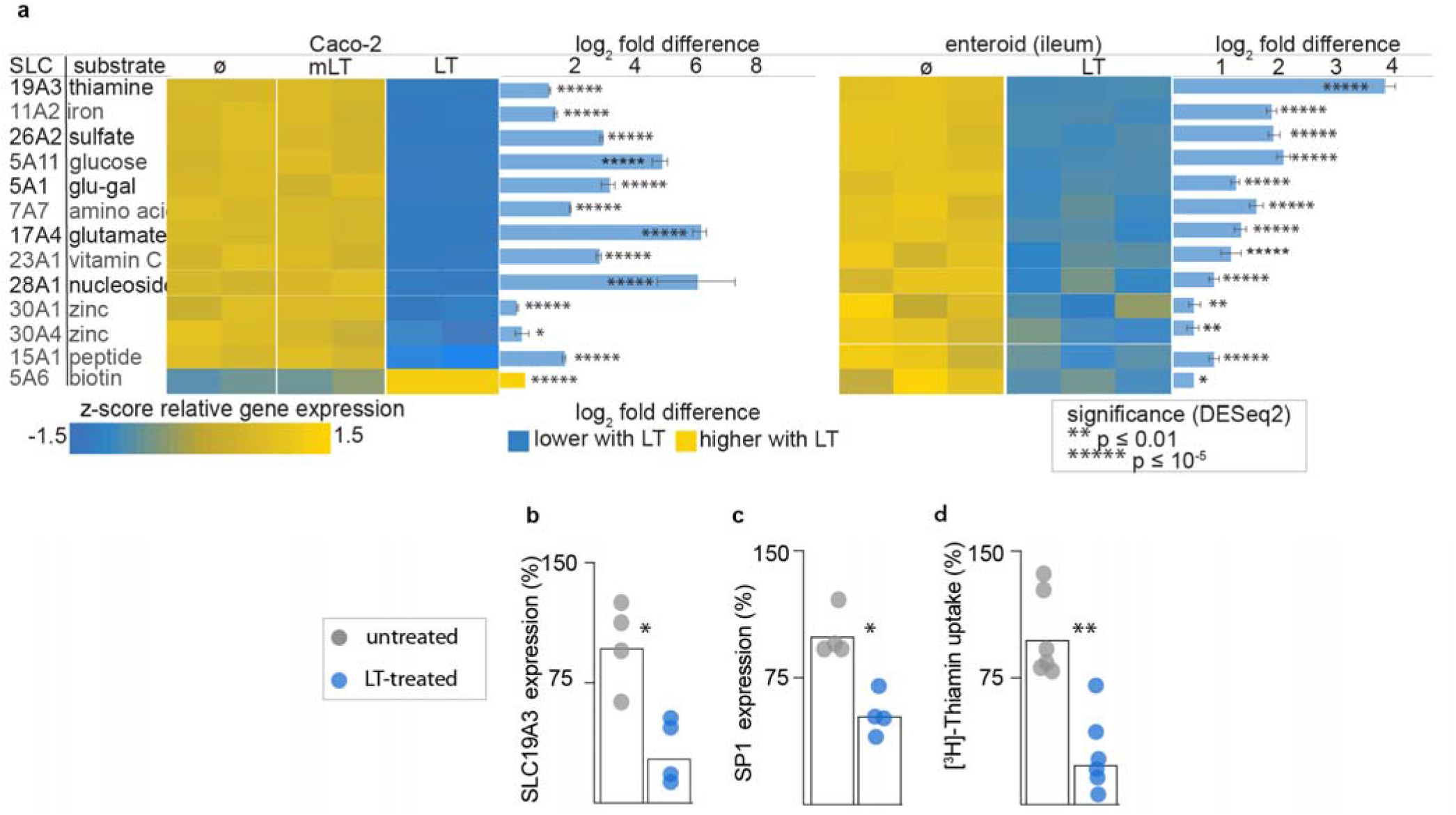
Heat-labile toxin alters the transcription of multiple brush border SLC genes. **a.** heat-map indicating key SLC genes modulated by heat labile toxin (LT) compared to enzymatically inactive E112K LT mutant (mLT), or untreated (ø) Caco-2 cells (left) and human small intestinal (ileal) enteroids (Hu235D, right). Real-time qRT-PCR confirming LT-mediated modulation of genes in ileal (Hu235D) enteroids encoding (**b.**) the major thiamine transporter SLC19A3 and (**c**) the SP1 cis regulatory element. Data reflect two independent experiments with 2 replicates each. **d.** uptake of [^3^H]-thiamine by Hu235 D cells is impaired following LT treatment. Data presented are from two independent experiments with n=3 replicates each. Bars indicate absolute fold change values + SE (* <0.05, **<0.01 Mann-Whitney nonparametric comparisons).

### LT-producing ETEC disrupt the absorptive architecture of the small intestine

To further study the potential impact of ETEC toxins on the intestinal architecture, we performed challenge studies in infant mice. Compared to sham-challenged (PBS) controls, or mice challenged with a toxin deficient strain of ETEC, we again noted down-regulation of genes involved in F-actin bundling, membrane cross-linking, and intermicrovillus adhesion complex formation, all required for intestinal microvilli (figure 5a) biogenesis. Likewise, on examination of small intestinal villi we found that production of villin in enterocyte brush borders was substantially decreased relative to sham challenged controls (figure 5b,c). In mice challenged with a wild type ETEC isolate that makes ST and LT, but not a LT/ST-toxin-negative mutant (<u>jf4763</u>, supplemental table 1), we observed significant alteration in architecture of the intestinal brush border with significant shortening and disorganization of the microvilli (figure 5d,e; supplemental figure 4a-d) reminiscent of the earlier ultrastructural studies of patients with tropical sprue ^17^ and *V. cholerae* infections^25^. Similarly, we found that in mice challenged with a strain containing an isogenic mutation in *eltA* (<u>jf571</u>, supplemental table 1) encoding the LT A subunit, which still makes heat-stable toxins, the microvillus architecture was preserved (figure 5f,g), suggesting that LT is the principal toxin underlying the enteropathic changes to the enterocyte surfaces.

**Figure 5.**
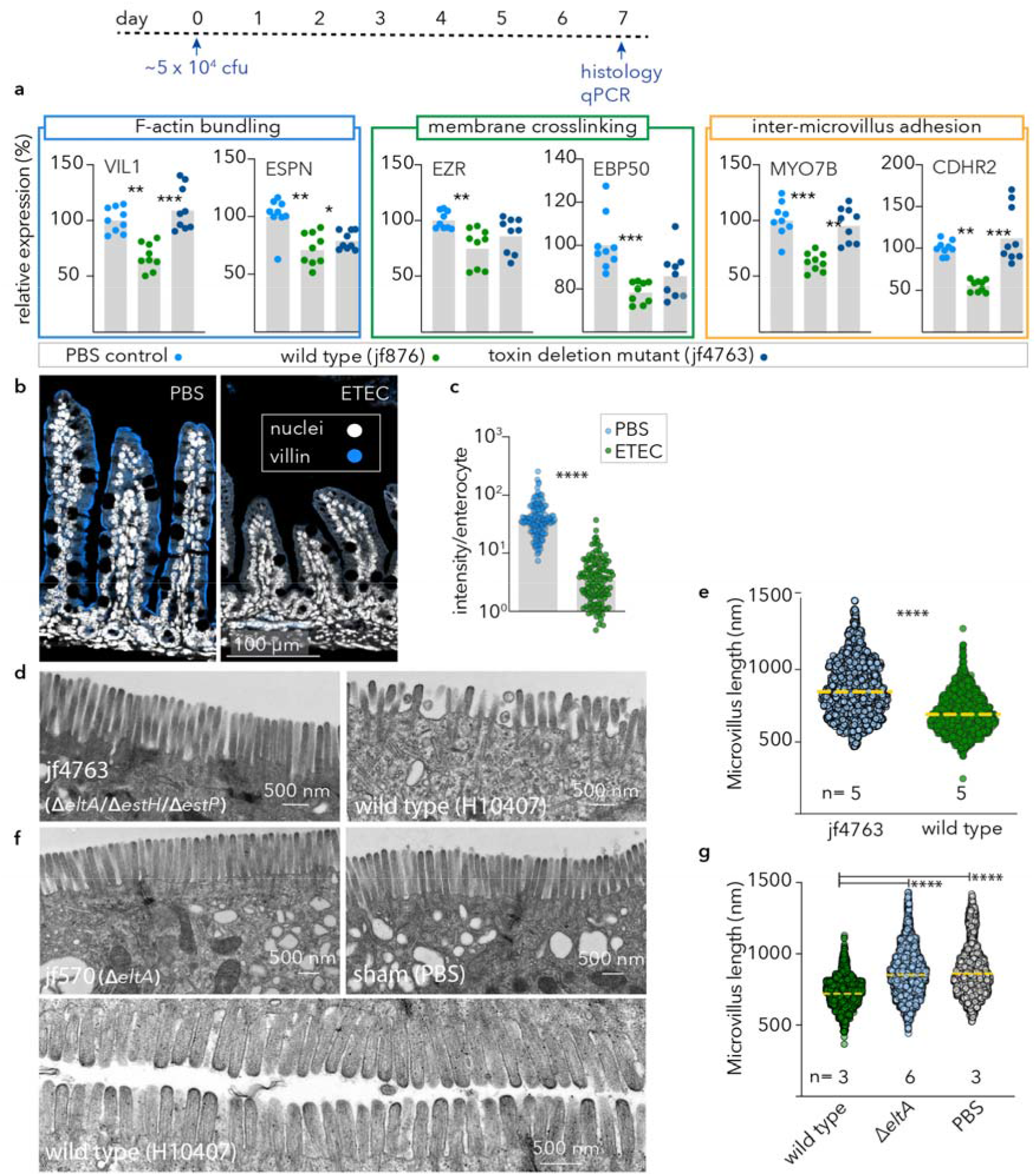
ETEC disrupts *in vivo* formation of small intestinal microvilli. Timeline at top depicts challenge with ETEC or control nontoxigenic isolate, or sham (PBS) challenge. **a.** Quantitative PCR results for genes involved in brush border development in small intestinal samples obtained from infant mice (n=9/group) 7 days after challenge with toxigenic ETEC (jf876), nontoxigenic ETEC (jf4763, LT^−^/ST^−^) PBS controls. Comparisons between data represent ANOVA, Kruskal-Wallis testing where ***p≤0.001, **p≤0.01, and *p≤0.05. **b.** Immunofluorescence images of small intestinal sections showing villin expression (blue), and nuclei (white). **c.** mean villin fluorescence intensity normalized per enterocyte (n=4mice/group). **d.** representative transmission electron microscopy (TEM) images of the small intestinal brush border from mice challenged with toxin-negative (Δ, left) and toxigenic ETEC (wt, right). **e.** microvillus length ****p≤0.0001 by Mann Whitney (two-tailed) nonparametric comparisons. Data represent geometric mean length from n=5 mice per group in three independent experiments. **f.** TEM images from mice challenged with jf570 (*eltA*::Km^R^), sham PBS controls, or mice challenged with wild type ETEC. **g**. length of microvilli (dashed horizontal lines represent geometric means). ****p<0.0001 by Kruskal-Wallis.

Despite the dramatic toxin-dependent changes to the absorptive surface of the intestine, the early growth kinetics of suckling mice challenged a single time with either wild type bacteria or a heat-labile toxin deletion mutant were surprisingly similar (supplemental figure 5A) and paralleled those of sham-challenged controls. However, enteropathy in young children is thought to reflect damage elicited by repeated infections. Children in endemic regions typically suffer *multiple* ETEC infections before their 2^nd^ birthday, and the risk of enteropathic sequelae increases multiplicatively per episode^8,42^. Therefore, to assess the contribution of repeated ETEC infections to growth impairment, we compared the growth kinetics of suckling mice challenged a single time to those repeatedly infected with wild type toxigenic ETEC. These studies demonstrated a clear impact of repeated infection on growth (supplemental figure 5B). Finally, we found that the growth kinetics of mice repeatedly challenged with wild type ETEC H10407 was significantly retarded relative to those challenged with the isogenic LT-mutant jf876 (supplemental figure 5C). Therefore, repeated infections in this model appear to recapitulate impacts observed following repeated ETEC infection in children, and our data suggest that these features are at least in part driven by LT.

### Maternal vaccination with LT prevents brush border disruption

To address whether enteropathic changes to the small intestine can be prevented by vaccination, and to further define the role of LT, we vaccinated mouse dams with heat-labile toxin and examined the brush border ultrastructure in suckling mice. Vaccinated dams but not sham vaccinated controls expressed significant levels of IgA and IgG in breast milk (figure 6a), consistent with increased levels of antibodies in the stomachs of infant mice (figure 6b). Notably, we found that maternal vaccination with LT completely abrogated changes to the microvilli (figure 6c,d) further substantiating the importance of LT in driving changes to the epithelial architecture.

**Figure 6.**
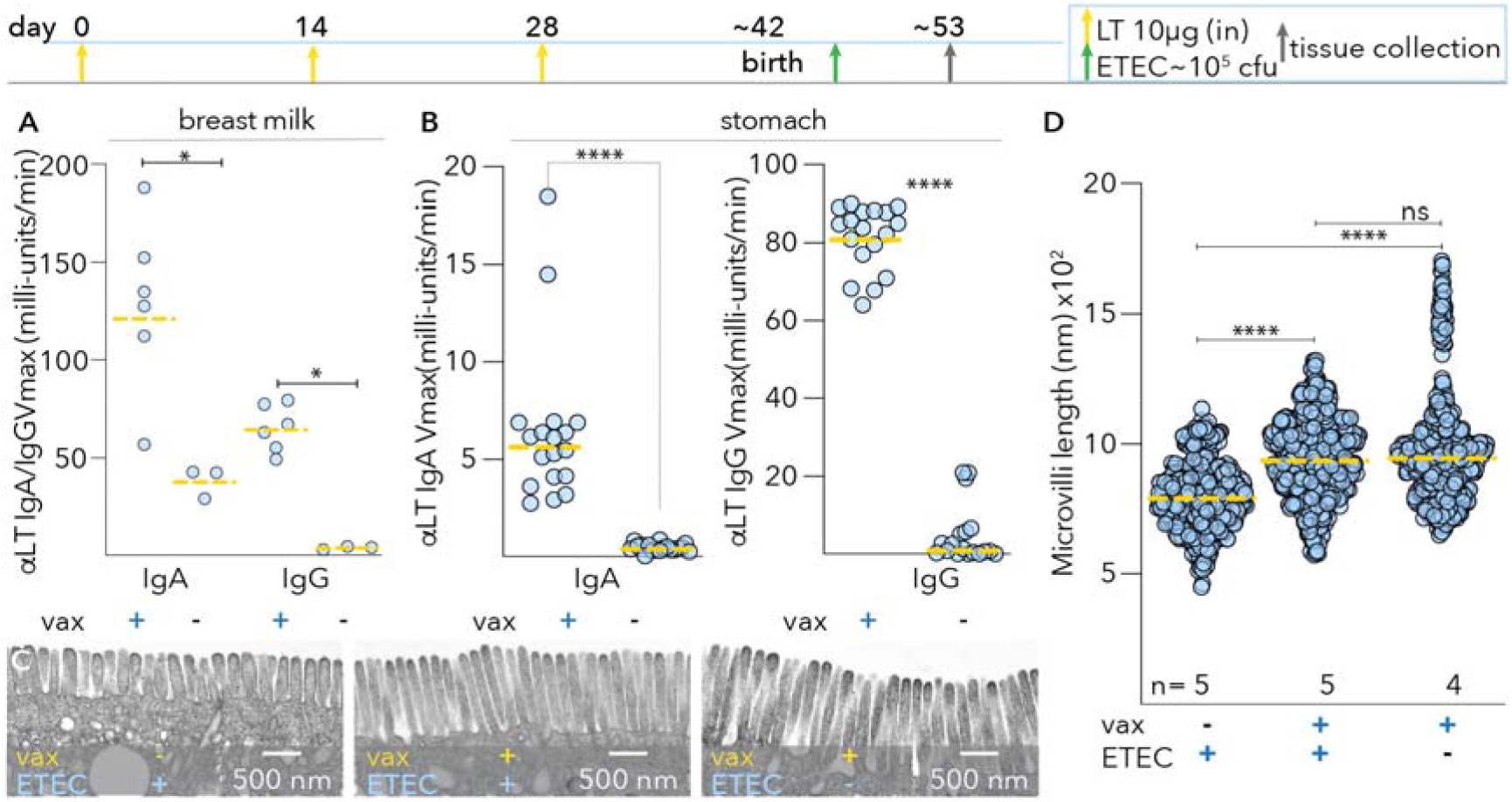
Maternal vaccination with LT mitigates microvillus disruption in neonatal mice. Timeline depicts vaccination and challenge (top): maternal intranasal (i.n.) vaccinations with 10 μg LT/immunization (yellow arrows on days 0, 14, 28) and neonatal challenge at 3 days of age (green arrow) followed by sacrifice and tissue collection at 7 days post infection (grey arrow). **A**. Kinetic ELISA data of breast milk anti-LT (IgA, and IgG in immunized dams and un-immunized controls). *<0.05 by Mann-Whitney two-tailed nonparametric testing. **B.** Anti-LT antibodies in the gastric contents of neonatal mice at day 53. **C.** Representative transmission electron microscopy images of brush border microvilli from unvaccinated mice challenged with wild type ETEC (left), vaccinated mice challenged with wild type ETEC, and vaccinated unchallenged controls. **D.** Microvillus lengths (based on image analysis of n=5 mice group) ****<0.0001 Kruskal-Wallis comparisons.

### LT modulates key transcription factors that govern enterocyte development

Despite the marked alteration in transcription mediated by LT the majority of genes critical for brush border development lacked conserved CRE sites ^23^. Therefore, we performed transcription factor target enrichment analysis^43,44^ to identify potential transcription factors responsible for differential regulation of genes significantly (P < 0.05) upregulated or downregulated by LT in both Caco-2 and enteroid RNA-seq datasets (supplemental table 4, <u>supplemental dataset 1</u>). The upregulated genes were most significantly (p = 4.2 x-10^−3^) linked to transcription factor targets of AP-1 encoded by the *c-jun* gene, previously shown to be regulated by cAMP^45^, and to be involved in intestinal epithelial repair^46^. Notably however, downregulated genes were most significantly enriched in targets for the HNF4α transcription factor or its intestine specific paralog HNF4γ (P = 1.9x-10^−4^) that were recently shown to regulate multiple genes required for brush border development^47,48^. Importantly, PKA has also been shown to phosphorylate HNF4 α at a consensus recognition site within the DNA binding domain, shared with HNF4 γ, inhibiting transcription^49^.

Of note, the transcription of both paralogs was found to be significantly depressed following exposure of intestinal epithelia to LT (figure 7A,B), as were levels of HNF4γ in nuclear fractions from toxin-treated cells (figure 7C), suggesting that activation of cAMP can interfere with transcription mediated by HNF4. To further examine the impact of LT on HNF4-mediated transcription we introduced a transcriptional reporter plasmid containing 6 tandem copies of the HNF4 transcriptional response element (5’-CAAAGGTCA-3’) linked to a human codon optimized *Gaussia princeps* luciferase into Caco-2 cells. These assays demonstrated that HNF4-mediated transcription was dramatically reduced in cells treated with LT (figure 7D). Similarly, we found that relative to nuclei of intestinal epithelia cells in ileal segments from sham-challenged control mice, those from ETEC-challenged mice exhibited significantly less HNFγ (figure 7E) further supporting a role for ETEC in modulating the production of this important transcription factor. Chen, *et al* described a “feed-forward regulatory module” essential to enterocyte differentiation in which HNF4 and SMAD4 transcription factors reciprocally activate each other’s transcription. As would be predicted from this model, we found that transcription of SMAD4 was also impaired by LT (supplemental figure 6A,B) leading us to speculate that LT mediated phosphorylation of HNF4 by PKA, interrupts this critical transcription module (supplemental figure 6C-E).

**Figure 7.**
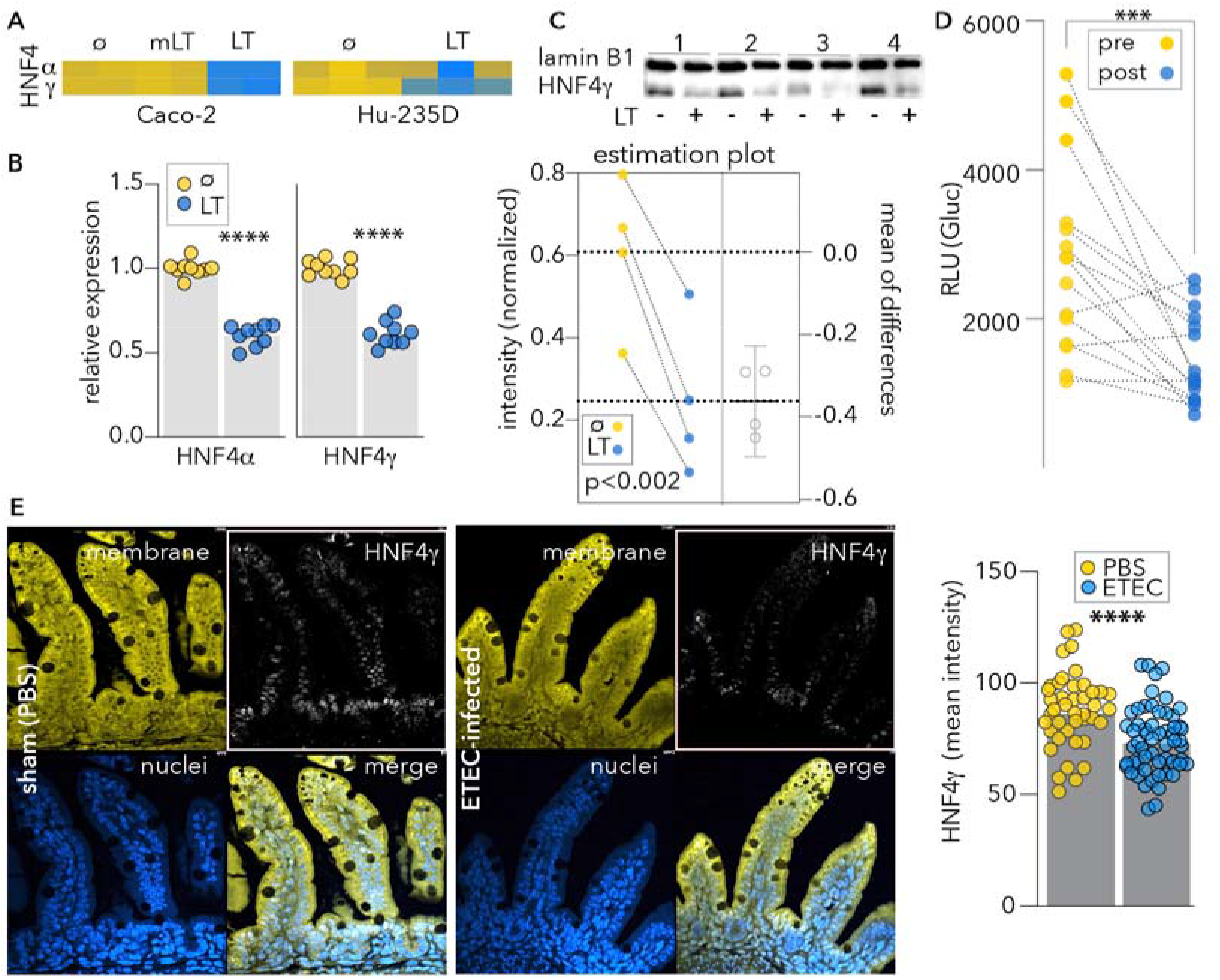
Enterotoxigenic E. coli heat-labile toxin impairs production of HNF4 nuclear receptors. **A.** heatmap demonstrating the impact of LT on transcription of paralogous transcription factors HNF4α and HNF4γ in Caco-2cells (left) and ileal enteroids, (Hu-235D, right) ø = untreated, mLT = mutant LT. **B.** qRT-PCR (TaqMan) data confirming decreased transcription of HNF4 transcription factors following treatment of enteroids with LT. **C**. HNF4γ is decreased in nuclear fractions obtained from small intestinal enteroids following treatment with LT. Shown in the HNF4γ immunoblot are samples from 4 independent experiments with graph below representing quantitation of signal intensity normalized to the lamin B1 nuclear protein (p=0.0017, paired t-test, one-tailed). **D.** HNF4 transcription *Gaussia* luciferase reporter assay showing decrease in signal following treatment of TR104-transfected Caco-2 cells with LT. 2 experimental replicates (n=15 samples total) *** p<0.001 Wilcoxon matched pairs, one-tailed). **E.** Confocal microscopy of representative Ileal sections from sham-challenged (PBS) left, and ETEC-infected mice (right). Immunofluorescence intensity of HNF4γ signal in sections from n=5 control mice, and n=6 ETEC-challenged mice. Membranes (yellow) were strained with CellMask orange (ThermoFisher <u>C10045</u>), nuclei (blue) stained with DAPI, and HNFγ immunostaining represented in white. Each symbol represents a microscopic region of interest. Bars represent geographic means (p<0.0001, Mann-Whitney two-tailed comparisons).

Collectively, the current studies demonstrate that in addition to their known canonical effects on ion transport that culminate in watery diarrhea, ETEC toxins can drive appreciable derangement of enterocyte architecture and function by interfering with key pathways in intestinal epithelia that govern the formation of mature enterocytes capable of effective nutrient absorption. These findings have important implications for our understanding and prevention of enteropathic conditions linked to ETEC.

## Discussion

Understanding the molecular events that lead to sequelae of undernutrition and growth faltering following ETEC infections may be key to the effective design of prevention strategies including vaccines^50^. Although the molecular mechanisms involved in the fluid and ion fluxes into the intestinal lumen leading to diarrhea are firmly established, only recently has evidence emerged to suggest that ETEC toxins may incite previously unappreciated changes in small intestinal epithelia^51,52^. The present studies, initiated to identify additional effects of LT, were prompted by an appreciation that cyclic nucleotides, particularly cAMP, govern a multitude of cellular pathways potentially resulting in collateral impacts that extend beyond the acute episodes of diarrhea. Consistent with this model, we found that exposure of intestinal cells to heat-labile toxin altered the transcription of hundreds of genes. A central theme highlighted in the analysis of these transcriptional alterations is that LT affects major classes of genes involved in biogenesis of microvilli and function of the intestinal brush border, the major site of nutrient uptake in the small intestine, potentially offering a direct molecular link to sequelae of malnutrition and impaired growth in children.

The potential clinical relevance of the observations reported here is highlighted by remarkably similar ultrastructural alteration of intestinal epithelial cells seen in small intestinal biopsies of patients with acute cholera^25^ and tropical sprue^17^. In both entities, the brush border was noted to be abnormal, with shortened, irregular microvilli. Importantly, however despite the structural and functional similarity between LT and CT, clinical cholera, unlike ETEC infections, has not been linked to enteropathy or attendant sequelae. Whether this relates to the repetitive nature of ETEC infections compared to the durable protective immunity that follows a single *V. cholerae* infection^53^ is not presently clear. Similarly, while tropical sprue remains a leading cause of malabsorption in regions where infectious diarrhea is prevalent^54–56^, the most debilitating forms of this illness have not typically followed isolated cases of traveler’s diarrhea, but occur in resident populations or expatriates^57^ repeatedly assailed by diarrhea while residing in endemic regions^58^. Likewise, our data also highlight the potential importance of repeated infections on the development of sequelae.

The negative impacts of LT on brush border architecture with commensurate reduction in surface area available for nutrient absorption, are compounded by the alteration of multiple SLC genes that encode transporters critical for uptake of essential vitamins and other molecules. Importantly, small intestinal biopsies obtained from Zambian children with enteropathy and refractory stunting exhibited similar changes in SLC gene expression profiles^59^. The decreased transcription of molecules required for intestinal zinc uptake, in cells treated with LT is particularly intriguing given the known aberrations in zinc absorption in children with enteropathy^35,60,61^, the possible contribution of zinc deficiency to enteropathic changes^62^, and the salutary effects of zinc in treatment of children with diarrhea^63,64^.

Further study will be needed to precisely delineate the role of ETEC LT and ST enterotoxins, alone and in combination, in driving enteropathic changes to the intestine and sequelae. ST-producing ETEC were most strongly associated with moderate to severe diarrhea in GEMS^3^, and follow-on studies of children enrolled in these studies have also linked infections with ETEC encoding heat-stable toxin to growth faltering^7^. However, our earlier analysis of a global collection of more than 1100 ETEC isolates, including those collected in GEMS, demonstrated that slightly more than half of all ST-encoding strains also encoded LT, and that roughly one third of the isolates overall encoded LT alone, ST-LT, or ST only^65^. While LT-producing ETEC have been specifically linked to malnutrition among children in Bangladesh^27^, and our *in vitro* and animal studies point to the potential importance of LT, additional effort will be needed to correlate toxin-induced morphologic and functional perturbation of the intestinal brush border with outcomes in children. Importantly, the long-term morbidity associated with ETEC infections does not appear to correlate with the severity of diarrhea as both mild illness^4^, and perhaps asymptomatic colonization may lead to growth faltering.

Further refinement of animal models that can faithfully recapitulate features of enteropathy are also needed. Indeed, conventional mice lack genes that could be required to reproduce the full effects of ETEC. For instance, each of the carcinoembryonic antigen cell adhesion molecules (CEACAMs) that are substantially upregulated on human small intestinal epithelia in response to LT, and which we have recently shown to play a critical role in ETEC interactions with human small intestine^51^, are absent in mice.

While the precise mechanism underlying LT-mediated modulation of genes required for microvillus biogenesis and absorptive function of the brush border is presently unclear, stimulation of adenylate cyclase invokes many cAMP-responsive nuclear factors that may serve either to activate or repress transcription. Genes implicated in development of microvilli are mostly devoid of consensus palindromic (TGACGTCA) or “half” (CGTCA) cAMP response element (CRE) sites within their promoter regions for direct modulation by CREB^66^, which typically is involved as a transcriptional activator, and both CREB and CREM can yield several alternatively spliced variants that may act as either activators or repressors^67^. cAMP second messaging also engages multiple signaling pathways converging at CREB^68^, and PKA can phosphorylate and modulate the activity of multiple transcription factors to act either as transcriptional activators or repressors, including SP1^69^. Notably, putative binding sites for HNF4, a cAMP-modulated transcription factor^49^ known to regulate genes needed for formation of microvilli^47,48^, were significantly enriched in the promotors of genes downregulated by LT. Both HNF4α and HNF4γ possess canonical PKA recognition sites within their DNA binding motifs, and PKA phosphorylation of these sites interrupts transcription^70^. HNF4 activates the transcription of SMAD4, and in turn SMAD4 activates transcription of HNF4^71^. Both transcription factors then engage genes needed for effective differentiation of stem cells to mature enterocytes^71^. Modulation of the activity of this transcription factor by LT would therefore be predicted to have a marked impact on pathways critical to intestinal epithelial homeostasis.

We should also note that increases in cellular cAMP can impact multiple cellular pathways independent of PKA. Included among these are pathways governed by a more recently discovered family of cellular cAMP-binding molecules, exchange proteins activated by cAMP (EPACs). EPACs appear to play critical roles as guanine exchange factors that regulate GTPase proteins^72^ and are in involved in complex signaling networks implicated in cell growth, differentiation, and morphogenesis^73,74^.

cAMP can also exert potent epigenetic influences on transcription. CREB-binding protein (CBP) possesses intrinsic histone acetyl-transferase (HAT) activity^75^, and can therefore modulate chromatin remodeling, enhancing access of transcription factors. In addition, cAMP messaging through PKA leads to phosphorylation-dependent activation of the histone demethylase enzyme PHF2 to promote the transcription of multiple genes that can impact transition from stem cells to epithelial cells^76^.

Altogether it seems likely that multiple pathways governed by increases in cellular cAMP may underlie the morphologic and functional disruption of the brush border epithelial observed in our studies. Nevertheless, the data presented here provide compelling evidence that the heat-labile toxin of ETEC ultimately impacts multiple genes required for the biogenesis and function of the brush border, the major site of nutrient absorption in the human small intestine. The findings may have significant implications for our understanding of sequelae linked to ETEC including environmental enteropathy in young children, and tropical sprue in adults. The increased acknowledgement of long-term morbidity linked to ETEC and an improved understanding of the role of ETEC enterotoxins as drivers of this morbidity may also strengthen the case for vaccines^50^ specifically engineered to prevent both the acute illness and sequelae.

## Methods

### Culture and differentiation of Enteroids

Biopsy samples from adult patients undergoing routine endoscopy were obtained at Washington University School of Medicine after patient consent and approval from the Institutional review board.

Purified crypt cells from the ileum were re-suspended in Matrigel (BD Bisosciences, San Jose, California, USA) and 15 μL of re-suspended matrix gel was added to each well in 24 well plates. Plates were incubated at 37° C and 5% CO2 with 50% L-WRN conditioned media (CM) and 50% primary culture medium (Advanced DMEM/F12, Invitrogen) supplemented with 20% FBS, 2 mM L-glutamine, 100 units/mL penicillin, 0.1 mg/mL streptomycin, 10 μM Y-27632 (ROCK inhibitor, Tocris Bioscience, R&D systems, Minneapolis, MN, USA), and 10 μM SB431541 (TGFBR1 inhibitor, Tocris Bioscience, R&D systems).

To induce differentiation and polarization of enteroids, cells were washed once to remove Matrigel, followed by trypsinization and centrifugation at 1100 x g for 5 minutes. Cells were then re-suspended in 1:1 CM and primary medium with Y-27632 and SB431541 as described above. Cells were then plated on semiporous filters (Transwells^®^, 6.5 mm insert, 24 well plate, 0.4 μm polyester membrane, Corning Incorporated, Kennebunk, ME, USA) that had been previously coated with Collagen IV (Millipore Sigma). Transwell^®^ inserts were rinsed with DMEM/F12 with Hepes, 10% FBS, L-glutamine, Penicillin, and streptomycin. Cells were allowed to grow to confluency in 50% conditioned media (CM) and then changed to differentiation medium (5% CM in primary medium + ROCK inhibitor) for 48 hours. Differentiated cells were used for toxin treatment.

### Propagation of Caco-2 cells

Caco-2 intestinal epithelial cells were obtained from ATCC (ATCC HTB-37) and cultured at 37 °C, 5% CO_2_, in Eagle’s MEM supplemented with 20% of fetal bovine serum (FBS). To generate polarized monolayers, ~1 x10^5^ cells were seeded onto polystyrene membrane filters (0.4μM, 6.5mm diameter insert, Transwell, Corning) and grew for at least a week prior to toxin treatment. Media was replaced with fresh media every two days.

### Toxin treatment

Polarized differentiated cells were treated with heat-labile toxin (LT) in differentiation medium (100 ng/mL) in a volume of 700 μL at the basolateral side of the Transwell insert and 100 μL volume on the apical aspect of the monolayer. Treated cells were incubated at 37° C for 18 hours and then fixed for transmission electron microscopy or fluorescence microscopy, or lysed for RNA extraction (GE Healthcare, Buckinghamshire, UK). LT and mutant LT (E211K, mLT) were kindly provided by Dr. John D. Clements, Tulane University, New Orleans, Louisianna, USA.

### RNA extraction and cDNA synthesis

RNA was extracted using Illustra RNAspin Mini RNA extraction kit (GE Healthcare, Buckinghamshire, UK). Three biological replicates per treatment (untreated and LT treated) were submitted to Genome Technology Access Center (GTAC) at Washington University in St. Louis School of Medicine for RNA-seq library preparation. An Agilent Bioanalyzer was used to determine the integrity of RNA samples. cDNA was generated using Superscript™ Vilo™ cDNA synthesis kit (Invitrogen by Thermo Fisher) after normalizing RNA concentrations.

### RNA-seq Library Preparation

Library preparation was performed with 10 ng of total RNA, integrity was determined using an Agilent bioanalyzer. ds-cDNA was prepared using the SMARTer Ultra Low RNA kit for Illumina Sequencing (Clontech) per manufacturer’s protocol. cDNA was fragmented using a Covaris E220 sonicator using peak incident power 18, duty factor 20%, cycles/burst 50, time 120 seconds. cDNA was blunt ended, had an A base added to the 3’ends, and then had Illumina sequencing adapters ligated to the ends. Ligated fragments were then amplified for 12-15 cycles using primers incorporating unique index tags. Fragments were sequenced on an Illumina HiSeq-2500 using single reads extending 50 bases.

### RNA-seq data processing and analysis

RNA-seq reads were aligned to the Ensembl top-level human genome assembly with STAR version 2.7.3a. Gene counts were derived from the number of uniquely aligned unambiguous reads by Subread:featureCount version 1.54.1. Read counts were used as input for DESeq2 differential gene expression analysis (version 1.24.0)^77^ with default settings, and a minimum *P*-value significance threshold of 10^−5^ (after False Discovery Rate [FDR^78^] correction for the number of tests). Fragments per kilobase per million reads mapped (FPKM) values for relative gene expression were calculated from DESeq2-normalized read counts, and Z-scores were calculated per gene using the average and standard deviations of FPKM values across samples. Log2 fold changes were identified from DESeq2 differential expression output. Pathway enrichment analysis for KEGG^79^ and Gene Ontology (GO)^80^ pathways among gene sets of interest was performed using the over-representation analysis tool provided on the WebGestalt^43^ web server (version 2019). Heatmap and bar graph visualization was performed with Microsoft Excel.

Transcript counts were produced by Sailfish version 0.6.3. Sequencing performance was assessed for total number of aligned reads, total number of uniquely aligned reads, genes and transcripts detected, ribosomal fraction known junction saturation and read distribution over known gene models with RSeQC version 2.3.

All gene-level and transcript counts were then imported into the R/Bioconductor package EdgeR and TMM normalization size factors were calculated to adjust for samples for differences in library size. Ribosomal features as well as any feature not expressed in at least the smallest condition size minus one sample were excluded from further analysis and TMM size factors were recalculated to created effective TMM size factors. The TMM size factors and the matrix of counts were then imported into R/Bioconductor package Limma and weighted likelihoods based on the observed mean-variance relationship of every gene/transcript and sample were then calculated for all samples with the voomWithQualityWeights function. Performance of the samples was assessed with a spearman correlation matrix and multi-dimensional scaling plots. Gene/transcript performance was assessed with plots of residual standard deviation of every gene to their average log-count with a robustly fitted trend line of the residuals. Generalized linear models were then created to test for gene/transcript level differential expression. Differentially expressed genes and transcripts were then filtered for FDR adjusted p-values less than or equal to 0.05.

The biological interpretation of the large set of features found in the Limma results were then elucidated for global transcriptomic changes in known Gene Ontology (GO) and KEGG terms with the R/Bioconductor packages GAGE and Pathview. Briefly, GAGE measures for perturbations in GO or KEGG terms based on changes in observed log2 fold-changes for the genes within that term versus the background log2 fold-changes observed across features not contained in the respective term as reported by Limma. For GO terms with an adjusted statistical significance of FDR <= 0.05, heatmaps were automatically generated for each respective term to show how genes co-vary or co-express across the term in relation to a given biological process or molecular function. In the case of KEGG curated signaling and metabolism pathways, Pathview was used to generate annotated pathway maps of any perturbed pathway with an unadjusted statistical significance of p-value <= 0.05. Genes significantly modulated in both Caco-2 cells and enteroids were subjected to further ontology-free analysis via CompBio v1.4 (PercayAI, Inc., www.percayai.com/compbio)^81^ to identify unifying biological themes in sets of genes differentially expressed between pairwise comparisons of different groups.

### Quantitative real time PCR

Quantitative real time PCR was performed using a QuantStudio 3 real-time detection system (Applied Biosystems). Fast SYBR Green master mix (Applied Biosystems / Thermo Fisher) was used for qPCR analysis. Disassociation curve analysis was performed to assess the specificity of amplification for each sample, and PCR product size was verified by agarose gel electrophoresis. Percent expression was normalized by GAPDH and analyzed using the comparative threshold cycle (Ct) method. Amplification of SLC19A3 transcripts from small intestinal enteroids was performed as recently described^82^ with relative gene expression normalized to β actin. HNF4a, HNFg, and SMAD4 gene expression was determined by TaqMan (ThermoFisher) using validated primer and probe sets. Primers and TaqMan probes used in this study are listed in Supplemental Table 2.

### Transcriptional reporter assays

To assess the impact of LT on HNF4-mediated transcription, pTR104 (GeneCopoeia) carrying 6 copies of the HNF4 transcriptional response element upstream of a secreted *Gaussia* luciferase (Gluc) reporter gene was transfected (<u>FuGENE, Promega</u>) into confluent Caco-2 cells. Media (EMEM, ATCC) were exchanged 24 hours after transfection, and after an additional 24 hours baseline samples of media were removed from each well and stored at −80°C. Wells were then treated with heat labile toxin (100 ng/ml) overnight (16 hours). At the end of treatment cell culture supernatants media were transferred to black walled microplates (Greiner 655086) and processed as directed (Secrete-Pair Luminescence, GeneCopoeia, Inc. Rockville, MD), followed by luminescence detection (Synergy H1, BioTek). Data are expressed as Relative Light Units (RLU) before and after treatment with LT.

SMAD4-mediated transcription was assessed by transient transfection of Caco-2 cells with the SBE4-luc plasmid containing 4 copies of the SMAD4 transcriptional response element 5’-GTCTAGAC-3’^83^. Following treatment with overnight treatment with LT cells were resuspended in 80 μl of EMEM media and mixed with an equal volume of ONEGLO EX Reagent (Nano-Glo^®^ Dual-Luciferase^®^ Reporter Assay System, Promega) for 20 minutes on an orbital shaker at 300 rpm, then read on the luminometer as above.

### Immunoblotting

#### SLC19A3, SP1

Total protein was extracted from LT toxin-treated (100ng/ml) human differentiated enteroid monolayers (235D line) and untreated control using radioimmunoprecipitation assay (RIPA) buffer (Sigma) containing protease inhibitor cocktail. An equal amount (~25 μg) of the proteins were loaded on a NuPAGE 4-12% Bris–Tris gradient gels (Invitrogen) as previously described^82^, then blotted onto polyvinylidene difluoride (PVDF) membranes and probed with anti-SLC19A3 (1:1000; Cat# 13407-1-AP; Proteintech), or anti-SP1 (1:1000; Cat# ab124804; Abcam) antibodies and together with anti-beta actin (1:3000; Cat# sc-47778) primary antibodies. Specificity of the SLC19A3 antibodies was validated in our laboratory previously using different approaches that include overexpression of tagged protein or gene silencing^84^. Anti-SP1 antibodies were validated by the manufacturer using knockout cell lysate protein samples. The SLC19A3 and SP1 protein bands from the blot were then identified with corresponding anti-rabbit IR-800 dye (1:30,000; Cat# 926-32211; LI-COR Bioscience) and anti-mouse IR-680 dye (1:30,000; Cat# 926-68020; LI-COR Bioscience) secondary antibodies incubation for 1 h at room temperature. Relative expression of specific proteins was calculated by comparing the fluorescence intensities in an Odyssey infrared imaging system (LI-COR Bioscience) with respect to corresponding beta actin signal.

#### Villin

Differentiated LT-treated and control enteroid monolayers were lysed to using NE-PER™ nuclear and cytoplasmic extraction reagent (Thermo Scientific). Cell membrane pellets were solubilized in PBS containing 1% Triton X-100 supplemented with protease inhibitor cocktail (Pierce™ protease inhibitor mini, Thermo Scientific). Equal amounts of total protein were loaded on a 4-20% gradient SDS-PAGE gel (Mini Protean TGX, Bio-Rad), then blotted onto nitrocellulose membranes and probed with anti-villin mouse monoclonal antibody (1:1000; Cat# SC-66022; Santa Cruz Biotechnoogy) followed by detection with HRP-conjugated anti-mouse secondary antibody (1:1000; Cat# 7076S; Cell Signaling Technology). Blots were developed with Clarity Western ECL substrate (Bio-Rad) and imaged with c600 imaging system (Azure Biosystems). Relative Villin expression signals were then analyzed with respect to corresponding Coomassie stained gel using imageJ analysis software.

### Thiamin Uptake

Initial rates (30 min; at 37 °C) of carrier-mediated thiamin uptake were examined in LT-treated (100 ng/ml; overnight) and untreated control differentiated small intestinal enteroid monolayers incubated in Krebs–Ringer buffer (pH 7.4) containing [^3^H]-thiamin (15 nM). Enteroid monolayers were then washed with ice-cold Krebs–Ringer buffer followed by lysis with NaOH and neutralization with 10 N HCl. The radioactive content was counted using a liquid scintillation counter as described previously^82^. Uptake of thiamin by its respective and distinct carrier-mediated mechanism was determined by subtracting uptake of [^3^H]-thiamin in the presence of a high pharmacological concentration (1 mM) of unlabeled thiamin from uptake in their absence; all uptake data points were calculated relative to total protein content (in milligrams) of the different preparations and presented as percentage relative to simultaneously performed controls.

### Confocal microscopy

Cell monolayers were fixed with 2% paraformaldehyde for 30 min at 37 °C and then washed 3× with PBS prior to blocking with 1% BSA in PBS for 1 h at room temperature. Villin expression was detected using 1:100 dilution of anti-villin antibodies raised in mice (Santa Cruz Biotechnology) for 1 hour at room temperature followed by additional 1 hour incubation with 1:200 dilution of fluorescence-tagged goat anti-mouse IgG Alexa Fluor 488 (Thermofisher) (Supplemental Table 5). Cell membranes were stained with 1:2000 dilution of CellMask orange (Invitrogen) and nuclei were counterstained with DAPI (1:1000) and mounted on glass slides using Prolong Gold antifade reagent (Invitrogen). Images were captured and analyzed on a Nikon C2 confocal microscope equipped with NIS-Elements AR 5.11.01 software (Nikon).

### Electron Microscopy

For ultrastructural analyses, *in vitro* grown differentiated polarized monolayers of human ileal enteroid samples as well as mouse intestinal biopsy samples were fixed in 2% paraformaldehyde/2.5% glutaraldehyde (Ted Pella, Inc., Redding, CA) in 100 mM sodium cacodylate buffer, pH 7.2 for 2 hours at room temperature and then overnight at 4°C. Samples were washed in sodium cacodylate buffer and postfixed in 2% osmium tetroxide (Ted Pella, Inc) 1 hour at room temperature. Samples were then rinsed in dH20, dehydrated in a graded series of ethanol, and embedded in Eponate 12 resin (Ted Pella, Inc.). Sections of 95 nm were cut with a Leica Ultracut UCT ultramicrotome (Leica Microsystems Inc., Bannockburn, IL), stained with uranyl acetate and lead citrate, and viewed on a JEOL 1200 EX transmission electron microscope (JEOL USA Inc., Peabody, MA) equipped with an AMT 8 megapixel digital camera and AMT Image Capture Engine V602 software (Advanced Microscopy Techniques, Woburn, MA). Images were analyzed using ImageJ software for microvilli length and structures.

### Identification of potential transcription factor target sites

The CREB targets database^66^ (http://natural.salk.edu/CREB/) was used to identify potential categorical CRE full (TGACGTCA) or half (TGACG/CGTCA) sites within the promotor regions of genes modulated by heat-labile toxin. Following previously described protocol^23^, 5 kilobase upstream sequences from each human gene were retrieved from Ensembl^85^ and were searched for the CREB binding sites TGACGTCA and TGACGTAA (and their reverse complements), based on sequences retrieved from the TRANSFAC^86^ database (matrix ID: V$CREB_01). CREB sequences matches were identified in the upstream sequences of 1,302 genes. WebGestalt^87^ was used for overall functional enrichment of potential transcription factor binding sequences upstream from genes that were significantly (p<0.05) up- and down-regulated in both Caco-2 cells and enteroids.

### Suckling mouse challenge studies

Studies in mice were performed under protocol 20-0438 approved by the Washington University in Saint Louis Institutional Animal Care and Use Committee. In neonatal mouse challenge studies, 3-day-old CD-1 mice (male and female) were inoculated with wild type ETEC (H10407), isogenic toxin deletion mutants, or sterile PBS (sham negative controls). Inocula were prepared from frozen glycerol stocks maintained at −80°C grown overnight (~16 hours) in 2 ml Luria Bertani (LB) broth in a shaking incubator (at 37°C, 225 rpm). Cultures were then diluted 1:100 in 20 ml of fresh LB, grown for additional 2 hours, and bacteria harvested by centrifugation (6000 rpm, 5 minutes at 4°C). Pellets were then washed once in ice-cold sterile PBS, then resuspended to a density of ~1*10^5^ Colony forming units (CFU) per 20 μl inoculum. CFU in each inoculum were determined by plating serial dilutions LB-agar plates. Mice were then inoculated with ETEC directly into the stomach through the abdominal wall using a 31g insulin needle. Following infection, mice were marked with tattoo ink to define groups and returned to the cage. Seven days post infection mice were sacrificed, and the ileal tissue collected fixed for microscopy, or preserved for RNA extraction.

### Bacterial strain construction

Strains used in these studies are described in supplemental table 1. Strain jf4763 is devoid of all enterotoxins and was constructed from a previously engineered ST-negative mutant of H10407, jf2848^88^. Briefly, jf2848 was first transformed with pKD46, a lambda red recombinase helper plasmid with selection on ampicillin. Jf2848(pKD46) was then transformed with a PCR amplicon generated with primers _to replace the *eltAB* genes encoding LT with a spectinomycin resistance cassette resulting in jf4763. The strain was subsequently validated by PCR and by testing supernatants in GM-1 ganglioside ELISA to confirm lack of LT production.

### Vaccination and immunologic assessment

Adult female mouse dams were vaccinated intranasally three times at 2-week intervals with 10 μg LT holotoxin in PBS, with the last dose administered approximately 2 weeks prior to anticipated delivery. Control mice were mock vaccinated with PBS. Fecal pellets collected from vaccinated dams were used to assess mucosal responses to LT. To collect breast milk the dam was separated from the litter ~2hours prior to milking. Oxytoxin (<u>Sigma O3251</u>), 0.002 IU/g, was injected IP to stimulate milk production, followed by collection via pipette tip. Anti-LT immune responses in maternal fecal extracts and breast milk, and stomach contents and sera of neonates (male and female) were assessed by GM1 ganglioside^89^ kinetic ELISA^90^ for anti-LT IgA and IgG, using plates coated with GM1 (Sigma G7641), as previously described^91^.

## Data availability

RNA seq data were deposited at the Sequence Read Archive (SRA) on the NCBI website https://www.ncbi.nlm.nih.gov/sra under BioProject accession number PRJNA875141 (detail in supplementary table).

## Supplemental Data

### Supplemental Figures

**Supplemental figure 1.**
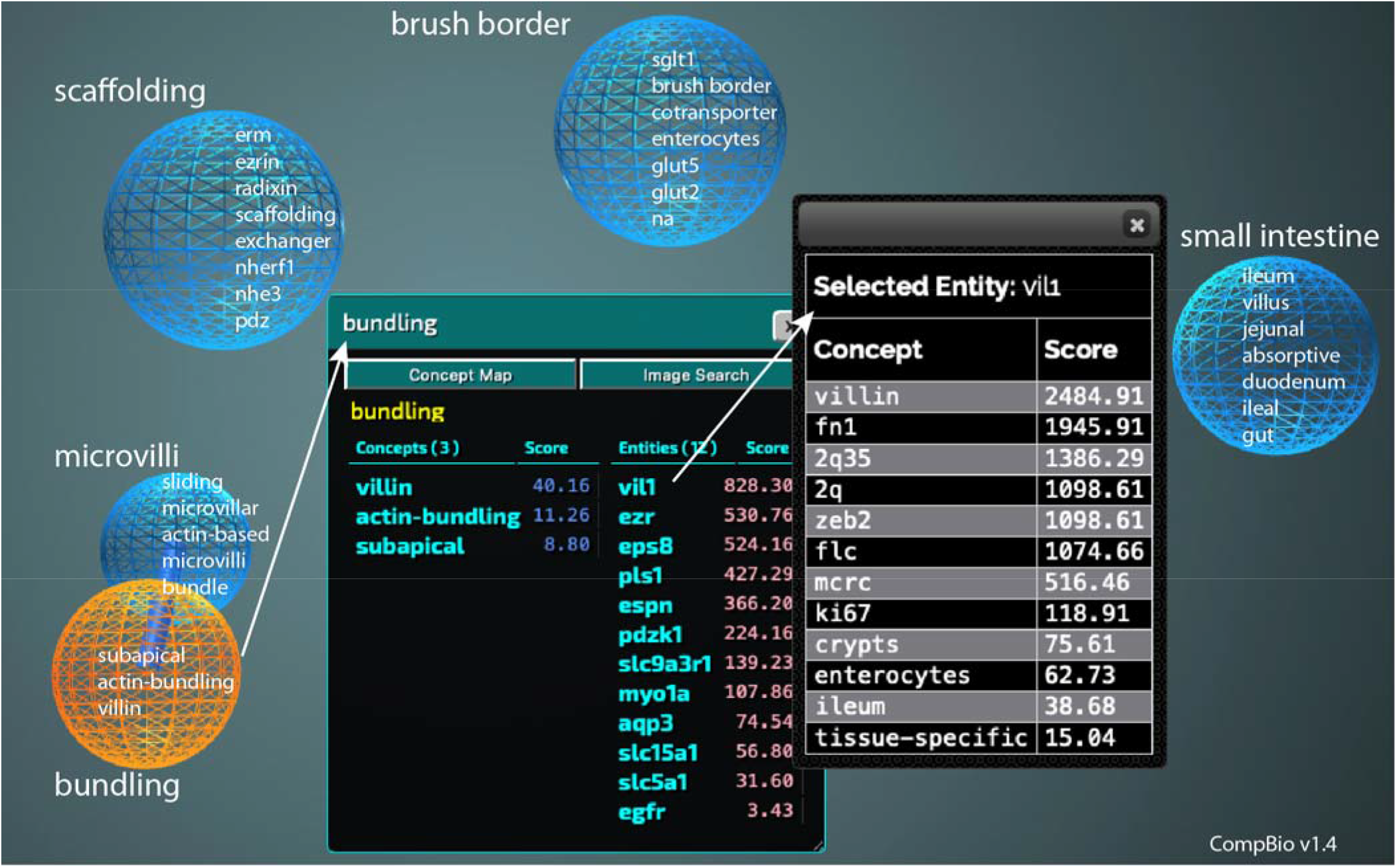
image from representative CombBio (v1.4) analysis of differential gene expression in LT-treated enteroids compared to untreated controls, highlighting themes (spheres) related to biogenesis and function of small intestinal microvilli. Concepts linked within each theme are listed on the corresponding sphere.

**Supplemental figure 2.**
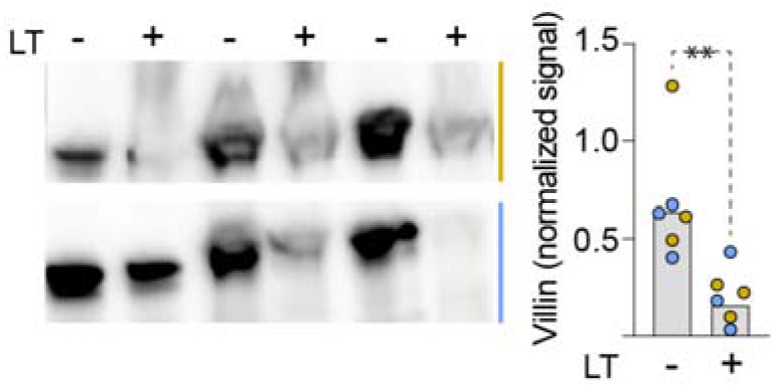
Villin production is decreased in LT-treated small intestinal enteroids. Shown are villin immunoblot signals in membrane preparations from 2 independent experiments (coded by color), each with 3 technical replicates. LT+ indicates overnight ~16 h treatment of ileal enteroids with heat-labile toxin [100 μg/ml]. Graph summarizes villin immunoblot signals normalized to total protein. (**p=0.0043 Mann Whitney, two-tailed comparisons).

**Supplemental figure 3.**
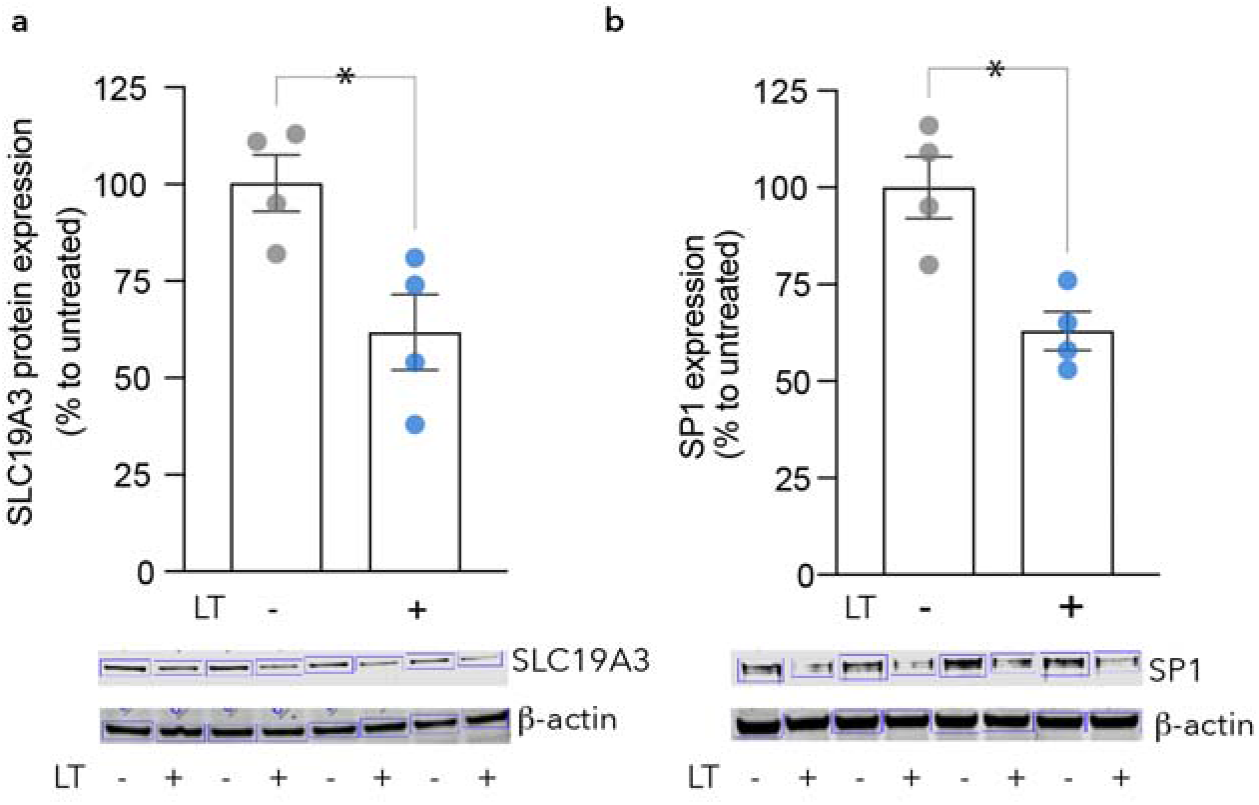
Impact of LT on expression of SLC19A3 and its associated SP1 transcription factor. Graphs depict summary of immunoblot band intensities of SLC19A3 (**a**) and the SP1 transcription factor (**b**) generated on probing LT-treated small intestinal enteroids (235D) relative to untreated controls. *<0.05 by Mann Whitney two-tailed analysis. Bars reflect geometric mean data ± SEM. The corresponding immunoblots from 4 independent experimental replicates are shown below each graph.

**Supplemental figure 4.**
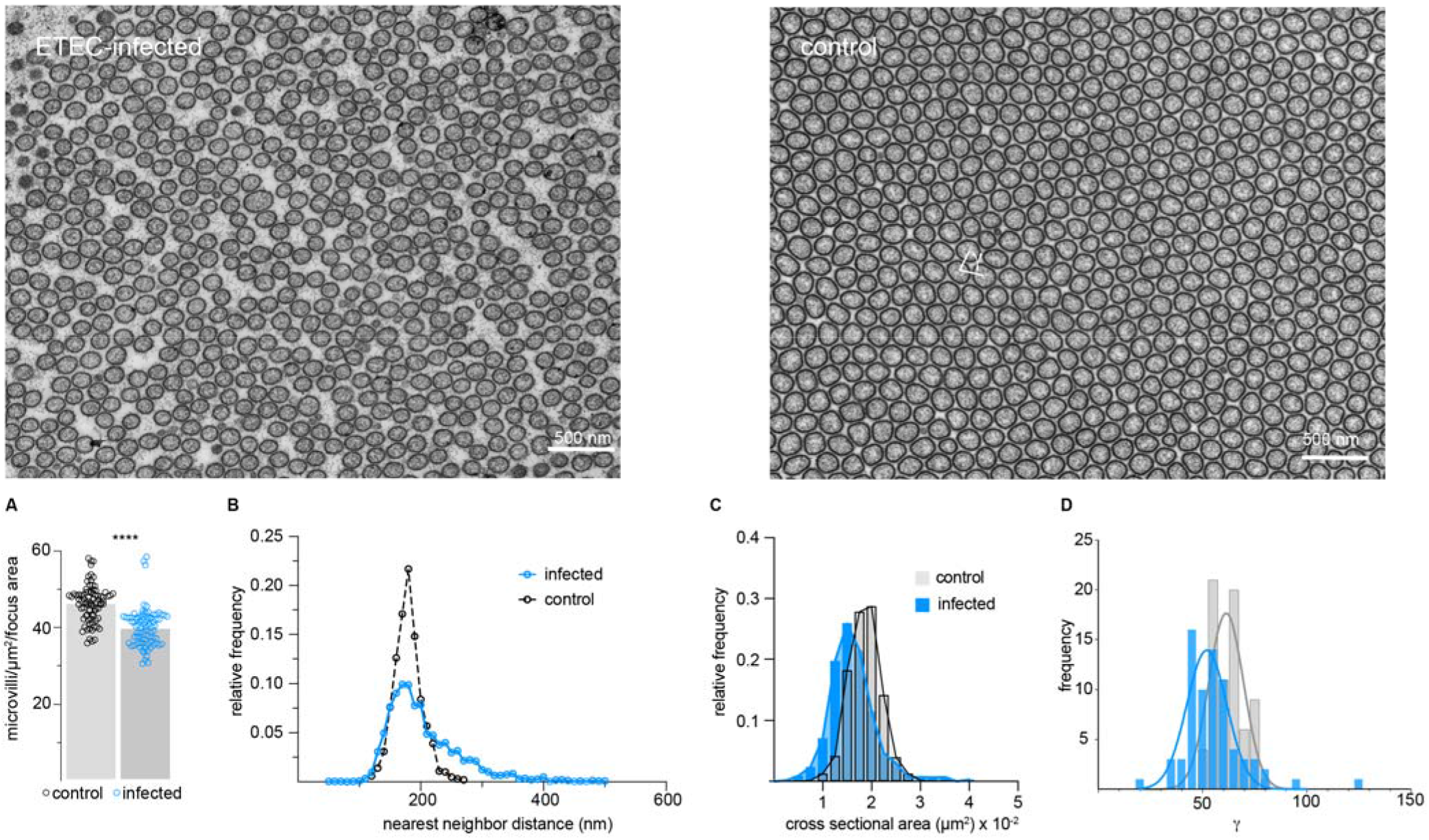
ETEC perturbs the ordered array structure of intestinal microvilli. Shown at top are representative TEM cross sections of infected and control mice (n=5) in each group. **A.** density of microvilli/μm^2^ (n=83 areas in infected and control). p<0.0001. **B.** distance between centers of adjacent microvilli (n=1001 control, n=2278 infected), p<0.0001 Mann-Whitney. **C.** cross sectional area of microvilli (n=550 control, n=1559 infected) p<0.0001. **D.** g angle distributions of microvilli from control and ETEC-infected mice. (analysis by Mann-Whitney, 2-tailed, nonparametric comparisons).

**Supplemental figure 5.**
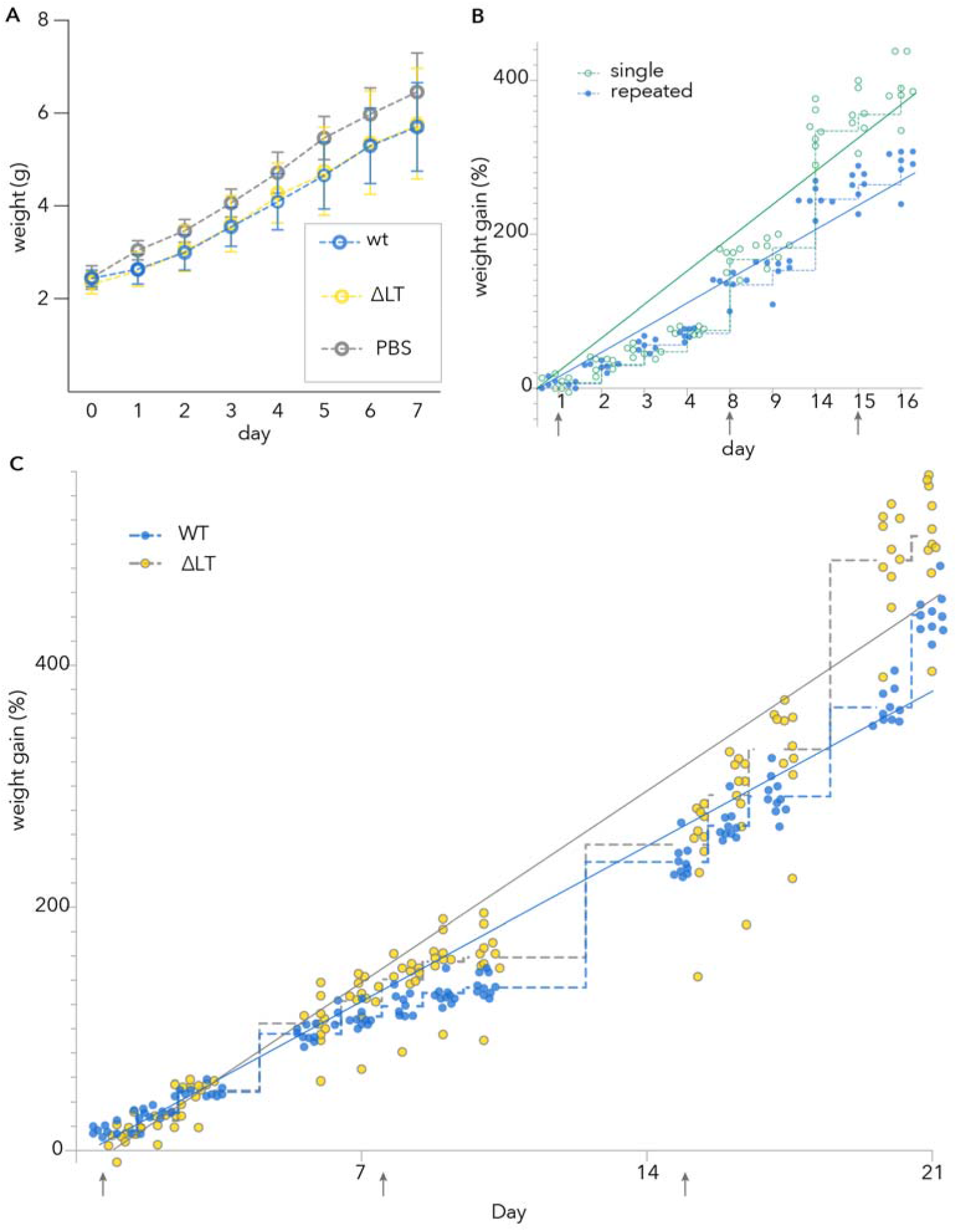
growth kinetics following challenge of suckling mice with ETEC. **A.** Short-term growth kinetics of mice singly challenged with wild type toxigenic ETEC H10407 (WT), LT mutant (Δ*eltA*, jf571), vs sham challenge (PBS) (n=24/group). data points reflect mean ± sd. **B.** Repeated vs single challenge of mice with wild type toxigenic ETEC H10407 (wt) (n=7 mice/group); dashed lines connect geometric mean values. Solid lines = linear regression. Comparison between singly and repeatedly infected by two-way ANOVA p=0.0004) **C.** LT contributes to growth impairment on repeated challenge. Shown are mice (n=10/group) repeatedly challenged with either H10407 or the LT deletion mutant jf571. Stepped lines connect geometric mean values. Solid lines indicate linear regression. (Comparison between WT and ΔLT by two-way ANOVA p=0.0078).

**Supplemental figure 6.**
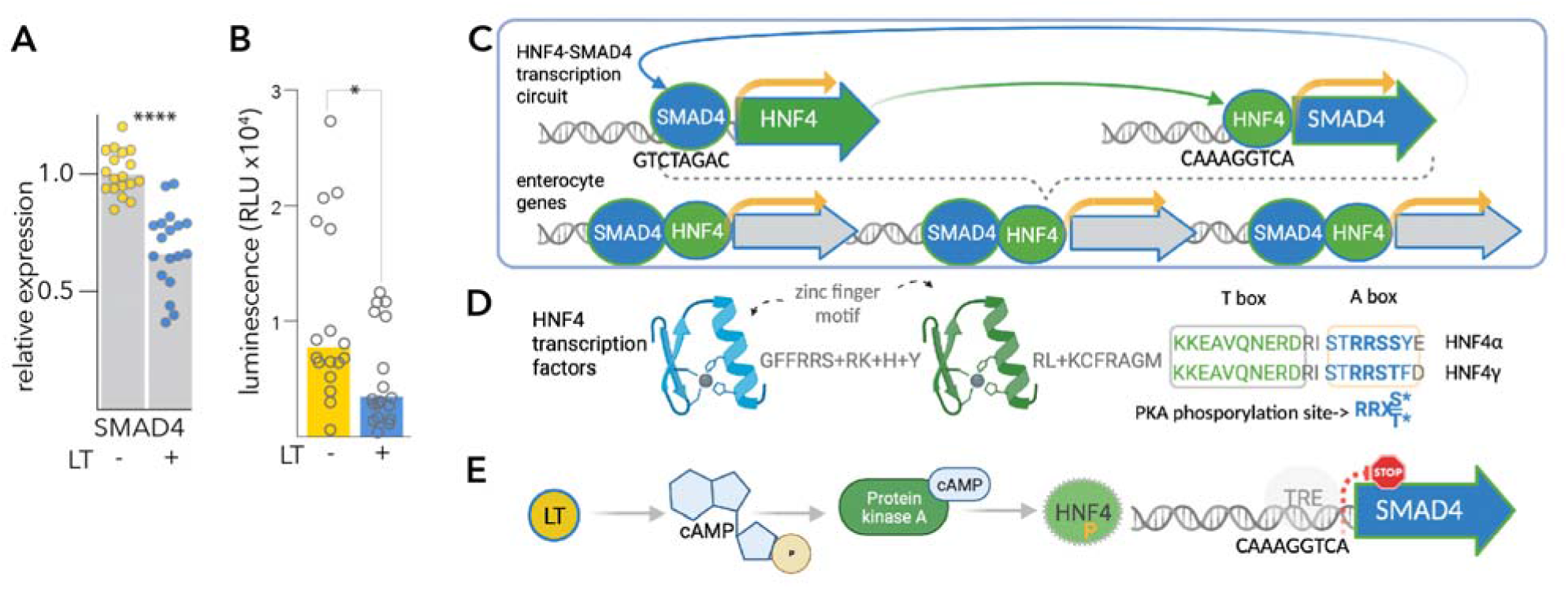
Potential mechanism underlying LT mediated disruption of brush border development. LT interrupts HNF4-SMAD4 mediated transcription. **A.** treatment of human-derived small intestinal enteroids (Hu235D) with LT leads to decreased SMAD4 transcription. Shown are TaqMan probe results from two independent experiments of polarized epithelial monolayers treated with LT. n=3 biological replicates x 3 technical replicates/ experiment =18 data points. ****p<0.0001 by Mann Whitney, two-tailed nonparametric comparisons to untreated cells. **B.** SMAD4-mediated transcription is decreased by LT. Shown are the combined results of three independent experiments in which Caco-2 cells transiently transfected with the SBE4-Luc plasmid containing 4 copies of the SMAD4 transcriptional response element were treated with LT (100 μg/ml x 16 hours). *=0.015 (Mann-Whitney). **C.** depiction of the HNF4-SMAD4 reciprocal transcription factor activation in which HNF4 activates transcription of SMAD4 and *vice versa* as (Chen, *et* al^71^). **D.** PKA-mediated phosphorylation of HNF4 paralogs interrupts a DNA-binding module (Viollet, *et al^70^*.). **E.** Summary of proposed mechanism by which LT leads to depletion of transcription factors essential for enterocyte differentiation. LT-mediated activation of PKA leads to phosphorylation of HNF4 rendering these transcription factors incapable of binding to *cis* transcriptional response elements (TRE) in promotors of target genes including SMAD4. Decreased SMAD4 transcription in turn depletes available HNF4.

### Supplemental Tables

**supplemental table 1.**
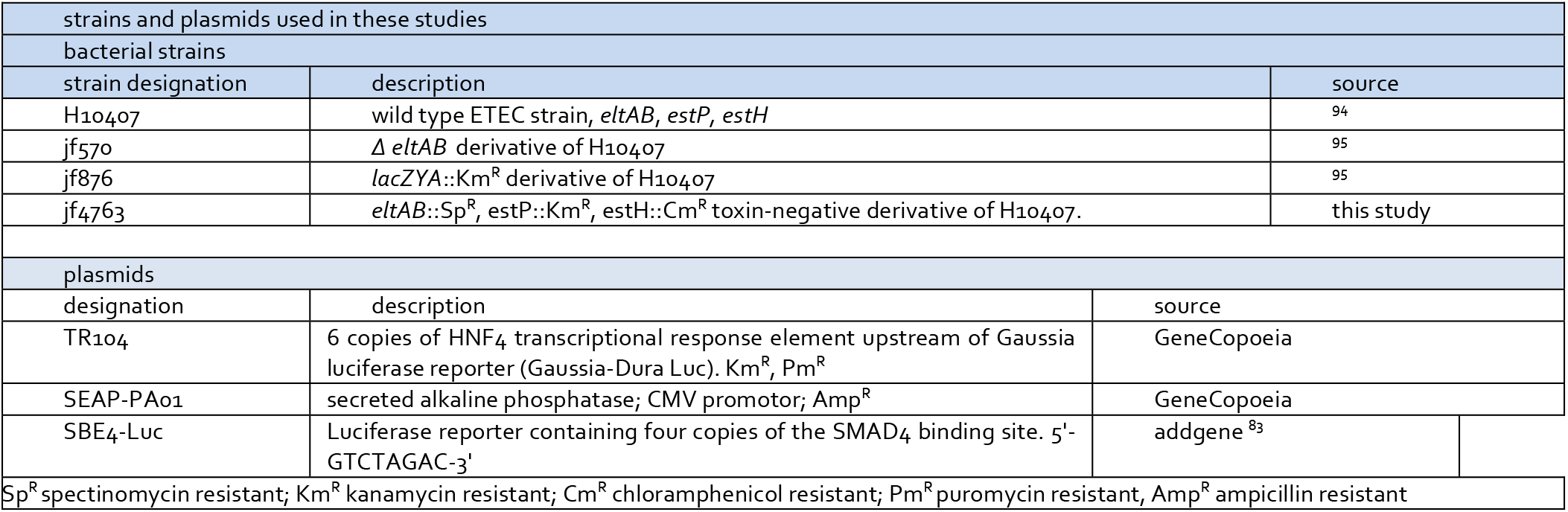

| strains and plasmids used in these studies |  |  |  |  |
| --- | --- | --- | --- | --- |
| bacterial strains |  |  |  |  |
| strain designation | description |  | source |  |
| H10407 | wild type ETEC strain, *eltAB*, *estP*, *estH* |  | 94 |  |
| jf570 | $\Delta$ *eltAB* derivative of H10407 | | 95 | |
| jf876 | *lacZYA::Km<sup>R</sup>* derivative of H10407 |  | 95 |  |
| jf4763 | *eltAB::Sp<sup>R</sup>*, *estP::Km<sup>R</sup>*, *estH::Cm<sup>R</sup>* toxin-negative derivative of H10407. |  | this study |  |
| plasmids |  |  |  |  |
| designation | description |  | source |  |
| TR104 | 6 copies of HNF4 transcriptional response element upstream of Gaussia luciferase reporter (Gaussia-Dura Luc). Km<sup>R</sup>, Pm<sup>R</sup> |  | GeneCopoeia |  |
| SEAP-PA01 | secreted alkaline phosphatase; CMV promotor; Amp<sup>R</sup> |  | GeneCopoeia |  |
| SBE4-Luc | Luciferase reporter containing four copies of the SMAD4 binding site. 5'-GTCTAGAC-3' |  | addgene <sup>83</sup> |  |
| Sp<sup>R</sup> spectinomycin resistant; Km<sup>R</sup> kanamycin resistant; Cm<sup>R</sup> chloramphenicol resistant; Pm<sup>R</sup> puromycin resistant, Amp<sup>R</sup> ampicillin resistant |  |  |  |  |

**supplemental table 2.**
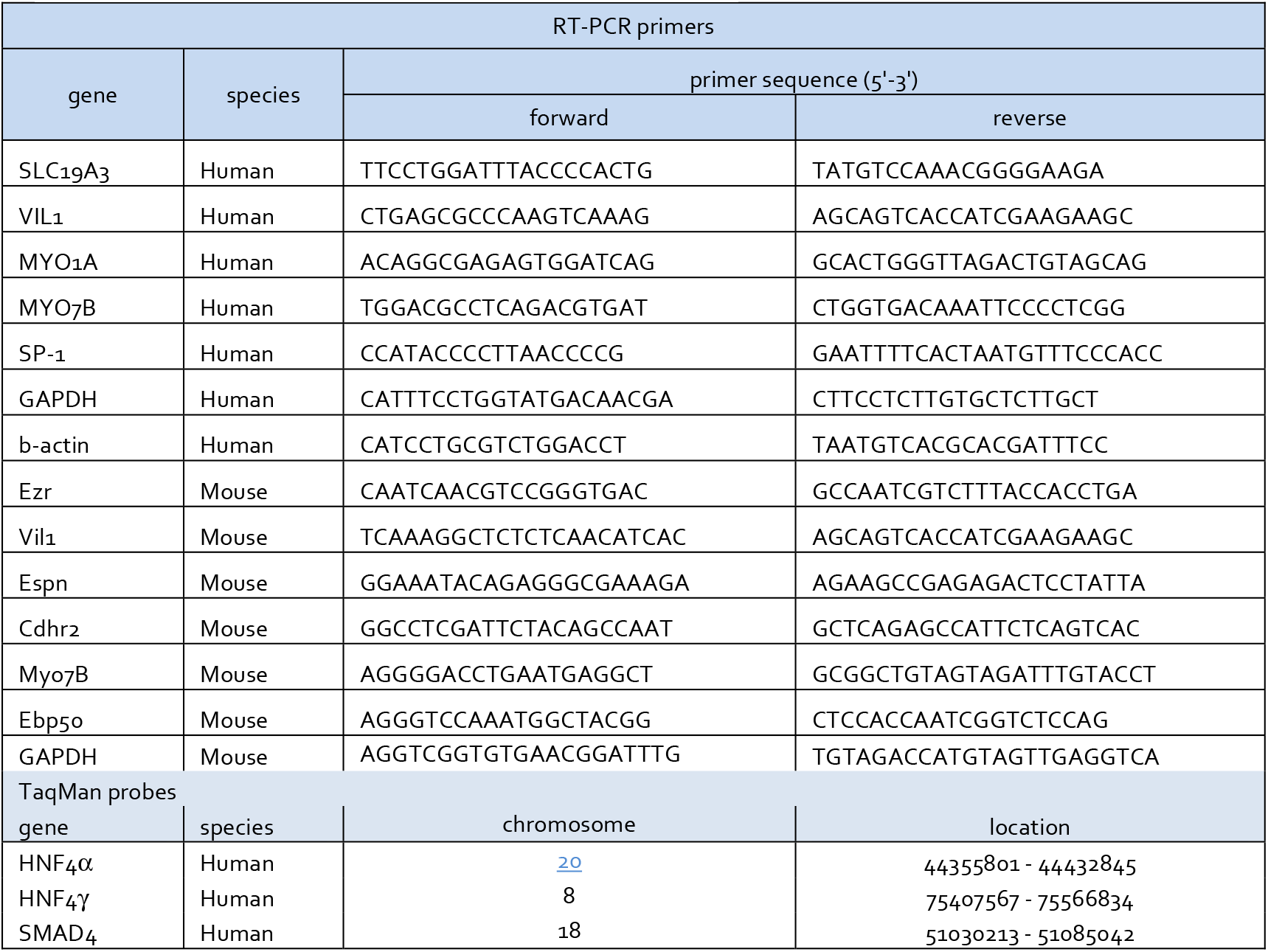
primers and probes for gene expression.

**supplemental table 3.**

| differential gene expression in gastrointestinal cells following treatment with LT |  |  |  |
| --- | --- | --- | --- |
| response to LT | dataset | genes (n)<br>p<0.05* | genes (n)<br>p<10<sup>-5</sup>* |
| higher | Caco-2 | 5,631 | 3,832 |
|  | ileal enteroids | 2138 | 746 |
|  | both<sup>†</sup> | 877 | 268 |
| lower | Caco-2 | 5,263 | 3,687 |
|  | ileal enteroids | 1,839 | 561 |
|  | both<sup>†</sup> | 1,013 | 311 |
| *differentially expressed genes identified by DESeq2<sup>93</sup><br><sup>†</sup> overlap in up- and downregulated genes was statistically higher than expected by random selection (p<10<sup>-10</sup>, binomial distribution testing)<br>Expression values of LT-treated enteroid cells are relative to untreated cells; LT-treated Caco-2 values are relative to untreated Caco-2 cells plus those treated with mLT. |  |  |  |

**supplemental table 4.**
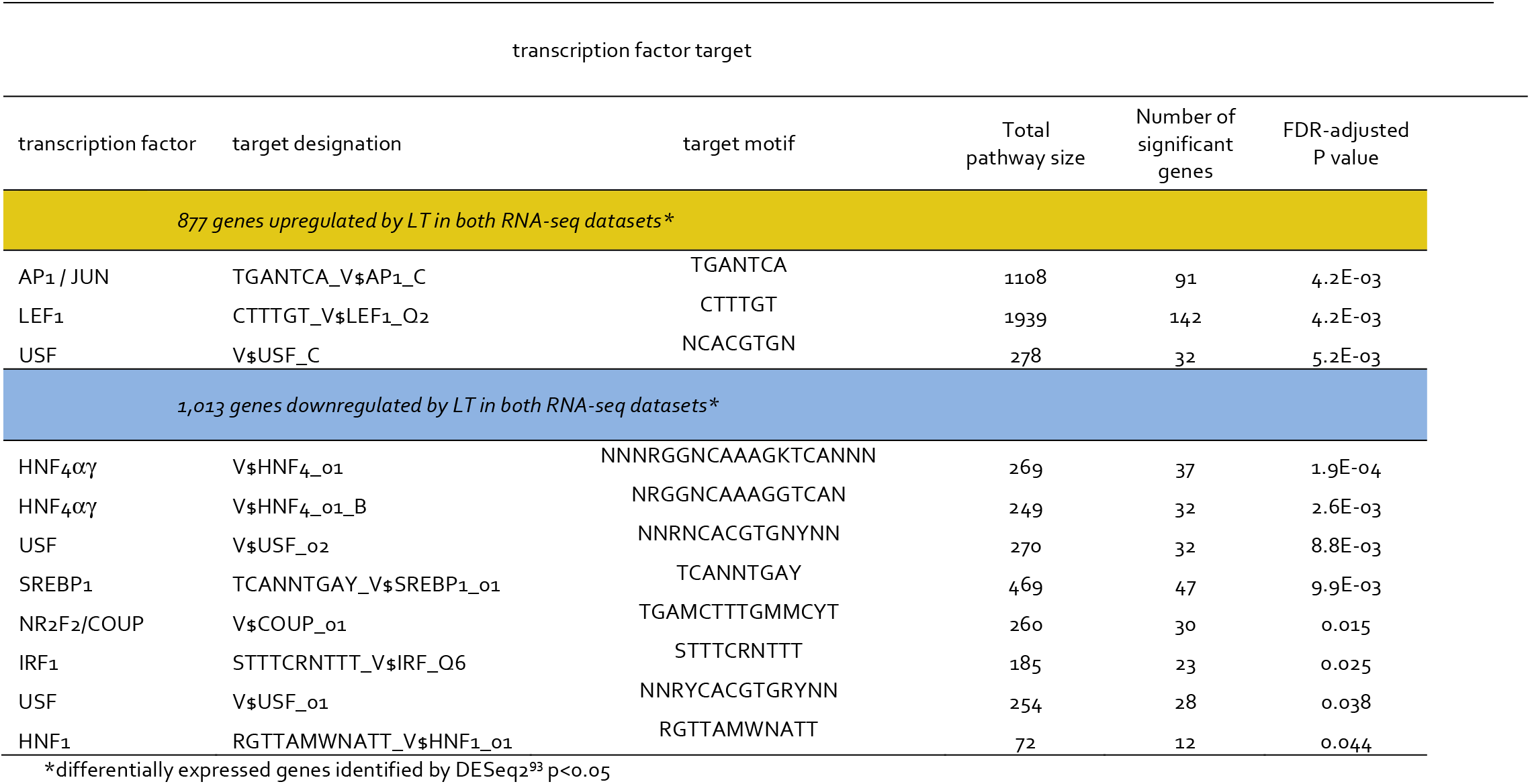
Transcription factor target sequence enrichment among genes differentially regulated in both RNA-seq datasets

**supplemental table 5.**

| antibodies used in these studies |  |  |
| --- | --- | --- |
| Specificity | description | source/reference |
| villin | Villin antibody (BDID2C3), mouse monoclonal IgG1 | Santa Cruz sc-66022 |
| mouse IgG | Goat anti-Mouse (H&L) IgG, highly cross-adsorbed antibody conjugated to Alexa Fluor 488 | ThermoFisher A-11029 |
| mouse IgA | Pre-absorbed, affinity purified goat anti-mouse IgA conjugated to HRP | Santa Cruz sc-3793 |
| mouse IgG | Affinity purified horse anti-mouse IgG conjugated to HRP | Cell Signaling 7076 |
| HNF4γ | rabbit polyclonal IgG against residues 327-408 of human HNF4γ | ThermoFisher 25801-1-AP |
| HNF4α | monoclonal IgG2a clone K9218) vs Baculovirus-expressed recombinant human HNF4α (aa 3-49) | ThermoFisher MA1-199 |
| SP-1 | rabbit monoclonal against synthetic peptide within Human SP1 aa 550-650. | abcam ab124804 |
| SLC19A3 | rabbit polyclonal IgG vs SLC19A3 | proteintech 13407-1-AP |
| β-actin | β-Actin (C4) is a mouse monoclonal antibody (IgG1 kappa light chain) raised against gizzard Actin of chicken origin. | Santa Cruz sc-47778 |
| rabbit IgG | IRDye<sup>®</sup> 800CW Goat anti-Rabbit IgG Secondary Antibody | LiCor 926-32211 |
| mouse IgG | IRDye<sup>®</sup> 680LT Goat anti-Mouse IgG Secondary Antibody | LiCor 926-68020 |

**supplemental table 6.**
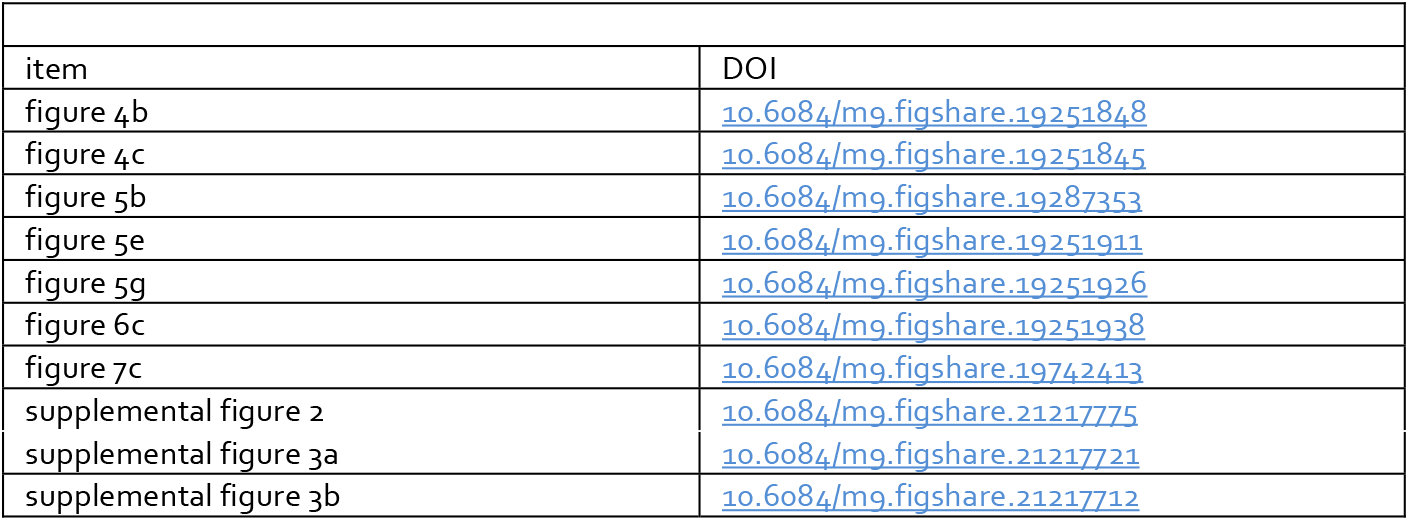

|  |  |
| --- | --- |
| links to original images |  |
| item | DOI |
| figure 4b | [10.6084/m9.figshare.19251848](https://doi.org/10.6084/m9.figshare.19251848) |
| figure 4c | [10.6084/m9.figshare.19251845](https://doi.org/10.6084/m9.figshare.19251845) |
| figure 5b | [10.6084/m9.figshare.19287353](https://doi.org/10.6084/m9.figshare.19287353) |
| figure 5e | [10.6084/m9.figshare.19251911](https://doi.org/10.6084/m9.figshare.19251911) |
| figure 5g | [10.6084/m9.figshare.19251926](https://doi.org/10.6084/m9.figshare.19251926) |
| figure 6c | [10.6084/m9.figshare.19251938](https://doi.org/10.6084/m9.figshare.19251938) |
| figure 7c | [10.6084/m9.figshare.19742413](https://doi.org/10.6084/m9.figshare.19742413) |
| supplemental figure 2 | [10.6084/m9.figshare.21217775](https://doi.org/10.6084/m9.figshare.21217775) |
| supplemental figure 3a | [10.6084/m9.figshare.21217721](https://doi.org/10.6084/m9.figshare.21217721) |
| supplemental figure 3b | [10.6084/m9.figshare.21217712](https://doi.org/10.6084/m9.figshare.21217712) |

## Acknowledgements

JMF was supported by funding from the National Institute of Allergy and Infectious Diseases (NIAID) of the National Institutes of Health (NIH) R01 AI126887, R01 AI089894, U01 AI095473; and the Department of Veterans Affairs (5I01BX001469-05). Research conducted by AS was also supported by National Institute of Allergy and Infectious Diseases of the National Institutes of Health under Award Number T32AI007172. We thank the Genome Technology Access Center in the Department of Genetics at Washington University School of Medicine for help with genomic analysis. The Center is partially supported by NCI Cancer Center Support Grant #P30 CA91842 to the Siteman Cancer Center and by ICTS/CTSA Grant# UL1TR002345 from the National Center for Research Resources (NCRR), a component of the National Institutes of Health (NIH), and NIH Roadmap for Medical Research. HMS was supported by funding from the Department of Veterans Affairs (I01BX001142) and the National Institutes of Health (DK5606 and AA018071). HMS is recipient of a Senior Research Career Scientist award # IK6BX006189.

The content is solely the responsibility of the authors and does not necessarily represent the official views of the National Institutes of Health, or the Department of Veterans Affairs.

